# Neuron-class specific responses govern adaptive remodeling of myelination in the neocortex

**DOI:** 10.1101/2020.08.23.263681

**Authors:** Sung Min Yang, Katrin Michel, Vahbiz Jokhi, Elly Nedivi, Paola Arlotta

## Abstract

Myelination plasticity plays a critical role in neurological function, including learning and memory. However, it is unknown whether this plasticity is enacted through uniform changes across all neuronal subtypes, or whether myelin dynamics vary between neuronal classes to enable fine-tuning of adaptive circuit responses. We performed *in vivo* two-photon imaging to investigate the dynamics of myelin sheaths along single axons of both excitatory callosal projection neurons and inhibitory parvalbumin^+^ interneurons in layer 2/3 of adult mouse visual cortex. We find that both neuron types show dynamic, homeostatic myelin remodeling under normal vision. However, monocular deprivation results in experience-dependent adaptive myelin remodeling only in parvalbumin^+^ interneurons, but not in callosal projection neurons. Monocular deprivation induces an initial increase in elongation events in myelin segments of parvalbumin^+^ interneurons, followed by a contraction phase affecting a separate cohort of segments. Sensory experience does not alter the generation rate of new myelinating oligodendrocytes, but can recruit pre-existing oligodendrocytes to generate new myelin sheaths. Parvalbumin^+^ interneurons also show a concomitant increase in axonal branch tip dynamics independent from myelination events. These findings suggest that adaptive myelination is part of a coordinated suite of circuit reconfiguration processes, and demonstrate that distinct classes of neocortical neurons individualize adaptive remodeling of their myelination profiles to diversify circuit tuning in response to sensory experience.

## Main Text

Myelin is a fundamental cellular structure in the vertebrate nervous system, and its precise formation and regulation are critical for complex neuronal function, including cognition ^1,2^. The importance of myelin for circuit function is made apparent by the severity of neurological diseases associated with its disruption ^3–5^. Given the impact of myelin on neuronal and network physiology, the question of differential regulation of myelination among the many diverse neuron classes in the central nervous system (CNS) is of fundamental importance.

In recent years, *in vivo* imaging studies have shown that myelination in the CNS continues through adulthood, along with constant remodeling ^6,7^ In addition, converging evidence suggests that myelin can be dynamically modulated by neuronal activity and contributes to nervous system plasticity throughout life ^8–11^. Myelin plasticity in response to experience helps to shape brain structure and function, including learning and memory ^5,12–14^. Previous studies of experience-dependent myelin plasticity have demonstrated the importance of myelination by newly-formed oligodendrocytes ^7,8,10–14^, but the neuronal substrates and dynamic properties of adaptively-remodeled myelin segments have not yet been addressed. It has been shown that different neuron subtypes have distinct patterns of axonal myelination, and, furthermore, that myelin profiles vary between individual cells within neuronal classes ^15–17^. A major unanswered question, therefore, is whether myelination plasticity affects different neuronal populations homogeneously, or whether adaptive remodeling has cell type-specific characteristics that may potentiate circuit tuning, either under normal conditions or driven by experience.

Here we investigate experience-dependent remodeling of myelination profiles on different classes of neurons by combining longitudinal *in vivo* two-photon imaging in the adult neocortex with genetic identification of cell types. Monocular deprivation (MD) is a classical model used to study sensory experience-dependent plasticity ^18^. In the adult mouse, MD induces a shift in response properties in both pyramidal neurons and inhibitory interneurons ^19^, but while interneurons extensively remodel both their dendritic and axonal compartments, layer 2/3 (L2/3) pyramidal neurons exhibit less remodeling ^20–23^. We therefore used MD in adult mice to interrogate whether experience might drive differential adaptive remodeling of myelination profiles on L2/3 callosal projection neurons (CPN) vs. parvalbumin-expressing interneurons (PV-INs). PV-INs in L2/3 make local connections to neighboring CPNs, thus participating in the same neural circuits ^24^, and are myelinated by the same pool of oligodendrocytes, including a subset of oligodendrocytes which myelinate both cell types simultaneously ^17^

We show that both excitatory CPNs and inhibitory PV-INs display remodeling of pre-existing myelin sheaths as well as *de novo* generation of myelin segments during normal conditions. However, monocular deprivation elicits an increase in myelin sheath dynamics only in PV-INs, and does not affect the myelination of CPNs nor the rate of generation of new myelinating oligodendrocytes. In addition, experience-dependent remodeling of myelination profiles on PV-INs coincides with an increase in axonal branch tip dynamics. These findings suggest that adaptive myelination is part of a complex and coordinated circuit-reconfiguration process which acts through differential effects on distinct neuronal subtypes.

### Homeostatic remodeling of pre-existing myelin

Previous reports have shown that activity and learning can modulate adult oligodendrogenesis as well as myelin dynamics ^7,11,25^. However, whether adaptive myelination is a cell type-specific process, perhaps allowing oligodendrocytes to differentially modulate the function of individual neuronal subpopulations, is still unclear. To study experience-dependent remodeling of myelination profiles and oligodendrocyte dynamics in the adult CNS, we used longitudinal dual-color *in vivo* two-photon imaging in the binocular area of the primary visual cortex (V1b) both during normal vision and through a period of ocular dominance plasticity induced by MD. Since > 90 % of the myelin in L2/3 of the neocortex wraps axons of either excitatory neurons (specifically, CPNs) or PV-expressing GABAergic interneurons ^16^, we compared myelin dynamics of these two functionally opposite neuronal classes (Figure 1A). We used the *Tbr2^CreERT2^; CAG^floxStop-tdTomato^* mouse line to specifically label L2/3 CPNs (Figure S1A-S1C), and the *PV^Cre^; cAG^floxStop-tdTomato^* mouse line to target PV-INs (Figure S1F and S1G). Both lines allow reliable labeling of their respective subpopulations as confirmed by immunohistochemistry (Figure S1). To generate the well-isolated labeling required for *in vivo* imaging, we optimized the tamoxifen induction protocol for the *Tbr2^CreERT2^; CAG^floxStop-tdTomato^* mouse line to sparsely label only the CPNs in the most superficial portion of L2/3 (Figure 1B, 1C, S1D and S1E).

**Figure 1.**
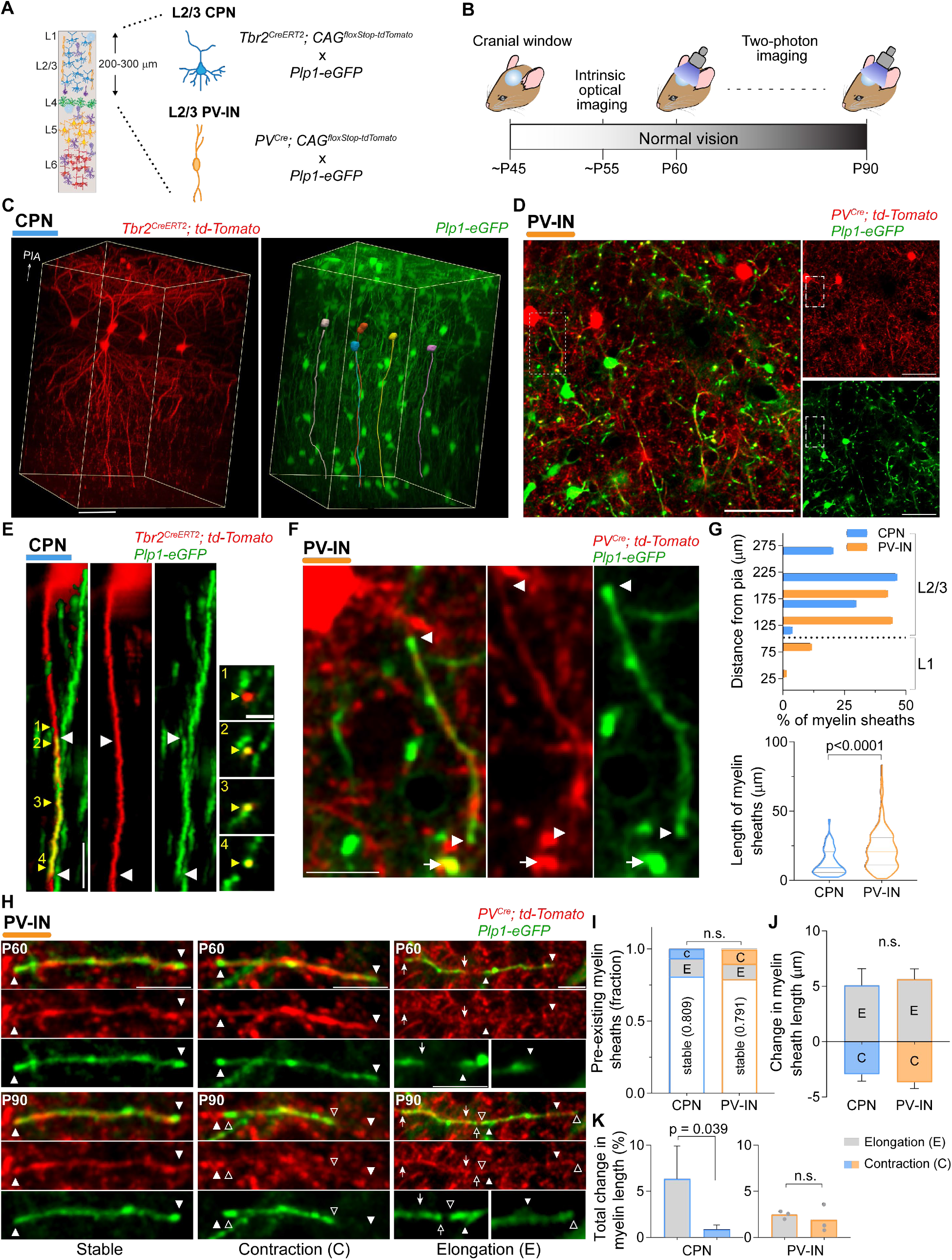
Pre-existing myelin sheaths on L2/3 PV-INs and CPNs present remodeling in adult mice. (**A**) Schematic of genetic labeling strategy. L2/3 CPNs and L2/3 PV-INs were fluorescently labelled in separate experiments using different Cre driver lines. (**B**) Experimental time course. Cells in V1b were imaged at P60 and P90. (**C**) Full imaged volume from a *Tbr2^Cre^* animal in three-dimensional perspective, showing CPNs (left) imaged simultaneously with oligodendrocytes and myelin (right). Skeletal reconstruction of cell bodies and primary axons of tdTomato^+^ CPNs are superimposed on the *Plp1-eGFP* image (right). (**D**) Representative single frame (~140 μm below pia) showing L2/3 PV-INs along with oligodendrocytes and myelin sheaths. (**E** and **F**) Representative images showing myelin sheaths (white arrowheads and arrows) on single axons of a superficial CPN [(E), three-dimensional image, yellow reconstruction in (C)] and a L2/3 PV-IN [(F), single frame, boxed area in (D)]. Cross-sections, at positions indicated by numbered yellow arrowheads, are displayed in the far-right column of (E). (**G**) Spatial distribution (top) and length (bottom, violin plot) of myelin sheaths at P90. Imaging of CPNs and PV-INs was performed 0-300 μm and 0-200 μm from the pia, respectively. (**H**) Maximum z projections (MZP) of myelin sheaths on PV-INs (arrows and arrowheads), showing stable (left, 3 frames), contracting (center, 3 frames), and elongating (right, 8 frames) internodes. (**I**) Number of dynamic (elongating and contracting) and stable myelin sheaths, normalized to the total number of pre-existing sheaths. (**J**) Length change of individual myelin sheaths. (**K**) Total change in myelin sheath length caused by elongations and contractions of pre-existing internodes, normalized to the total length of all myelin sheaths. Scale bars: 50 μm, (C) and (D); 10 μm, (E), (F) and (H); 5 μm, (E) inset. Data are mean ± s.e.m. For statistics, see table S1. n.s., not significant.

We combined this strategy for cell type-specific neuronal labeling with fluorescent detection of myelinating oligodendrocytes using the *Plp1-eGFP* transgenic line, allowing the simultaneous imaging of single axons from each neuronal type (tdTomato^+^), the myelin sheaths wrapping the axons (eGFP^+^), and the myelinating oligodendrocytes (Figure 1C, 1D and S2A). We confirmed that the eGFP signal from *Plp1-eGFP* mice faithfully reflected the presence and length of myelin sheaths by immunohistochemistry against myelin basic protein (MBP) and Cntnap1 (AKA Caspr), and by its spatial colocalization with single axonal branches from either CPNs or PV-INs (Figure 1E, 1F and S2B-S2E). We performed long-term *in vivo* two-photon imaging of layer 1-3 V1b in adult animals (postnatal day 60-90 [P60-90]) after the surgical placement of a cranial window and subsequent mapping of V1b using intrinsic optical signal (Figure 1A and 1B; see Materials and Methods).

We first performed two imaging sessions 30 days apart under normal vision to establish baseline changes in myelination. For PV-INs, we traced a total of 292 myelin sheaths, which were present at a density of 211 ± 17 myelin sheaths per 10^-2^ mm^3^ (n = 3 mice), consistent with the high rate of myelin coverage reported for this type of GABAergic interneuron ^16,17^ Cell bodies of PV-INs were distributed in L2/3 (10.7 ± 1.7 bodies), with no somata found in L1 (n = 3 mice); accordingly, 87.6 % of myelin sheaths on PV-INs were localized to L2/3, while only 12.4 % of them were in L1 (Figure 1G, top). Contrary to PV-INs, tracing of 47 myelin sheets on CPNs revealed that most of the CPNs were unmyelinated at P90 (Figure S3A-S3I), with only 20.5 % of this excitatory population possessing at least a single myelin sheath (Figure S3G). We also found that the onset of CPN myelination occurs late in development (~P30), and the rate of myelination is strikingly low but persistent throughout adulthood (4.5 % additional myelinated CPNs per month, from P30 to P210; Figure S3J-S3L).

We found that in mice with normal visual experience, the average myelin sheath was longer on PV-INs (22.4 ± 0.7 μm) than on CPNs (13.3 ± 1.3 μm) (Figure 1G, bottom), consistent with previous reports ^15,26^. Interestingly, pre-existing myelin sheaths displayed the capacity to elongate or contract over time (Figure 1H), with approximately 20% of the sheaths on both PV-INs and CPNs changing between P60 and P90 (Figure 1I). The fraction of dynamic segments and their change in length (range approximately −10 μm to +20 μm) is consistent with previous reports analyzing baseline, non-cell-type-specific myelin plasticity in S1 between P60 and P90 ^6^. For PV-INs, the number of elongation (*E*) and contraction (*C*) events was similar (*E*, 10.6 % vs *C*, 10.3 %), while pre-existing sheaths on CPNs showed a greater proportion of elongations (*E*, 12.7 % vs *C*, 6.4 %) (Figure 1I). Since the average change in sheath length is similar for elongations and contractions (Figure 1J), the overall remodeling of pre-existing myelin sheaths on PV-INs was balanced (*E*, 2.5 ± 0.2 % vs. *C*, 1.9 ± 0.8 %, of total myelin length; Wilcoxon matched-pairs signed rank test, p = 0.75, n = 3 mice), while there was a net increase in the level of myelination on CPNs (*E*, 6.3 ± 3.5 % vs. *C*, 0.9 ± 0.5 %, of total myelin length; Wilcoxon matched-pairs signed rank test, p = 0.039, n = 20 mice) (Figure 1K). Altogether, these data show that pre-existing myelin sheaths in the adult neocortex present neuronal cell-type specific patterns of homeostatic plasticity under normal vision, with L2/3 PV-INs displaying a balanced ratio of elongations and contractions, whereas CPNs exhibit shorter myelin sheaths and an overall elongation of segments over time. The unique myelin dynamics adopted by these distinct neuronal subtypes suggests that neuron type-specific mechanisms may implement the effects of myelination on circuit plasticity in the adult neocortex.

### *De novo* generated myelin sheaths are more dynamic

The cortex continues to add new myelin sheaths throughout life ^6,7,14^ To test whether new myelin segments have distinct dynamic characteristics compared to pre-existing internodes, we performed longitudinal *in vivo* two-photon imaging on mice with normal visual experience at one-week intervals between P60 and P90, and analyzed the pre-existing (identified in the first imaging session) and *de novo* (emerging during the course of the 4-week experiment) myelin sheaths on L2/3 PV-INs and CPNs (Figure 2A). Along with the longitudinal changes in pre-existing myelin sheaths described in the previous section, we found a high number of new myelin sheaths (~19.7 % of total internodes at P90; Figure 2B and 2C), on both CPNs and PV-INs. *De novo*-generated myelin sheaths had similar lengths as pre-existing internodes, such that the myelin length distributions across new and pre-existing sheaths were indistinguishable, for both neuronal populations (Figure 2D). In PV-INs, new myelin sheaths were continuously produced, with new sheaths identified in 15 out of the 16 weekly imaging sessions (Figure 2E and 2F) (the number of new sheaths in CPNs was not large enough to enable detailed analysis). The generation of new myelinating oligodendrocytes was rare (3 new oligodendrocytes across 4 mice), but in imaging sessions with new myelinating oligodendrocytes we saw an almost five-fold increase in both the rate of *de novo* generation of myelin segments and the total length of added myelin (Figure 2E and 2F).

**Figure 2.**
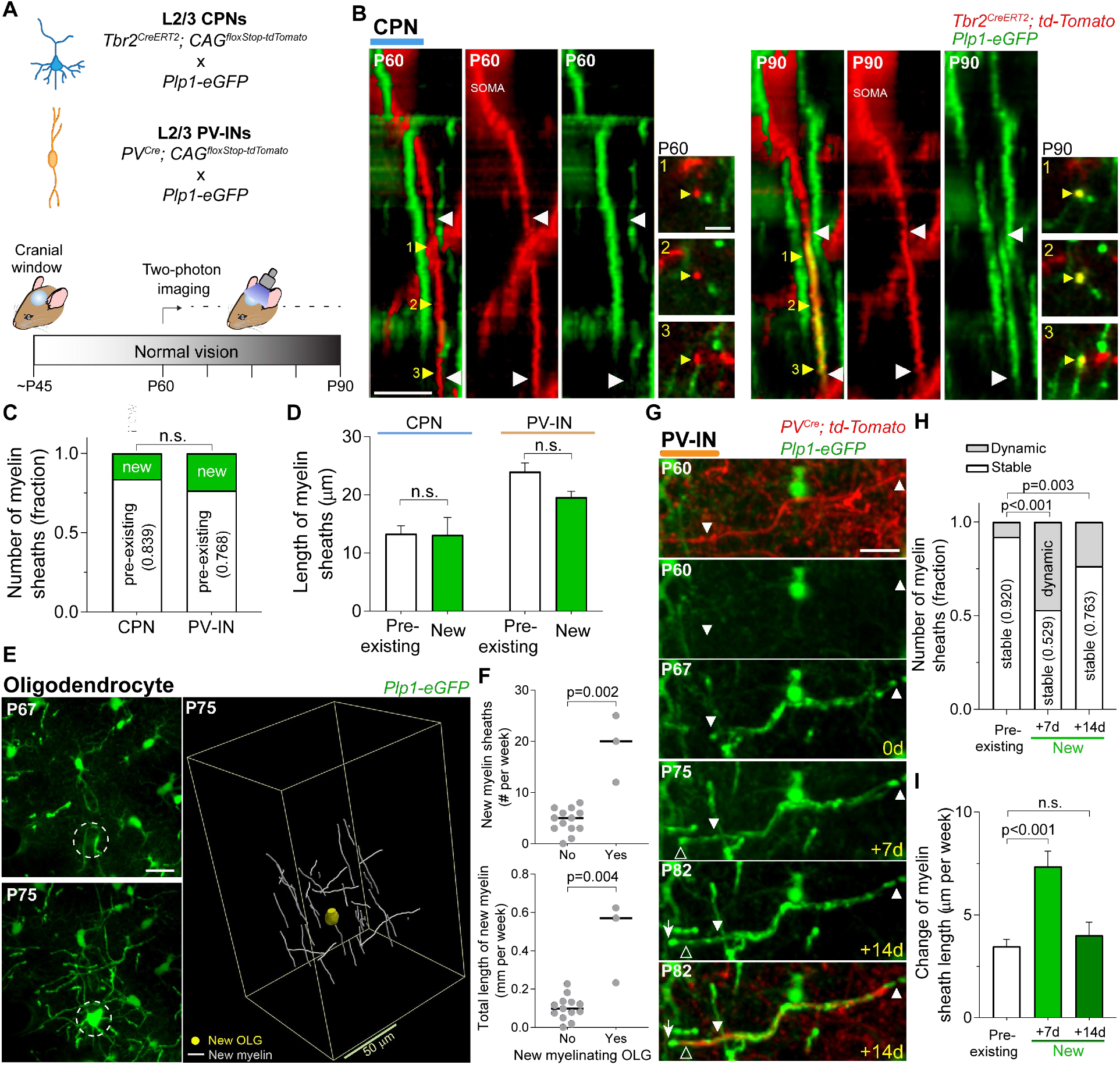
*De novo* generated myelin sheaths are more plastic than pre-existing internodes. (**A**) Schematic of cell and myelin labeling strategy (top) and experimental time course (bottom). Cells in V1b were imaged weekly between ~P60 and ~P90. (**B**) Representative three-dimensional images displaying a new myelin sheath (white arrowheads) generated between P60 and P90 on a single CPN axon (~150 μm from pia). Numbered yellow arrowheads point to the position of cross-section insets. (**C**) Fraction of pre-existing and *de novo* generated myelin sheaths at P90. (**D**) Length of individual myelin sheaths at P90. (**E**) Maximum z projections (MZP, 15 frames) showing the generation of a new myelinating oligodendrocyte (left, circle) and three-dimensional reconstruction of newly generated myelin sheaths on PV-INs, surrounding the cell body of the new oligodendrocyte (right; OLG, ~130 μm from pia). (**F**) Number (top) and total length (bottom) of new myelin sheaths on PV-INs, when a new myelinating oligodendrocyte was generated (Yes) or not generated (No) within the imaged volume. Each dot represents a single imaging session. (**G**) MZP (14 frames, ~115 μm from pia) displaying the remodeling of a *de novo* generated myelin sheath on a PV-IN (arrows and arrowheads). Timeline in yellow is relative to the first session in which we detected the new internode (P67 in the example). (**H** and **I**) Number of dynamic and stable myelin sheaths [(H), normalized to the total number of internodes in each group] and length change of internodes (I). We analyzed new internodes on PV-INs one and two weeks after they were first identified [see (G)] and compared them to the pre-existing sheaths. Scale bars: 10 μm, (B) and (G); 5 μm, (B) inset; 20 μm, (E). Data are mean ± s.e.m. For statistics, see table S1. n.s., not significant.

We next examined the level of remodeling of new sheaths compared to pre-existing sheaths, by comparing the imaging session in which individual new sheaths were first identified with sessions one and two weeks later (Figure 2G). It has been reported that newly-formed oligodendrocytes generate all of their myelin sheaths within 3 days of their differentiation ^25,27^, thus these imaging intervals extend well beyond the initial period of formation of new sheaths. While both new and pre-existing sheaths showed elongation and contraction, the new myelin segments exhibited a much larger ratio of elongations to contractions (54 *E*: 3 *C*, new sheaths; 31 *E*: 30 *C*, pre-existing sheaths). Notably, 47.1 % of the new myelin sheaths displayed changes in length during the first week after they were first detected, while only 8.0 % of pre-existing sheaths were remodeling during the same period (Figure 2H). The change in length of individual sheaths was also two times larger for new sheaths compared to pre-existing myelin (Figure 2I). Moreover, the remodeling rate of new sheaths remained higher than that of pre-existing internodes for at least two weeks after their generation. Taken together, these results demonstrate that, while rarer, new myelin sheaths are more dynamic than pre-existing myelin, and greatly contribute to myelin plasticity in adult mice.

### Experience-dependent myelin remodeling is neuron type-specific

It is unclear whether experience-dependent reconfiguration of neuronal circuits is accompanied by a remodeling of myelination profiles in a homogenous or a cell type-specific fashion. To address this question, we compared the changes in myelin dynamics and the generation rate of new myelinating oligodendrocytes before and during monocular deprivation. MD drives ocular dominance plasticity in the binocular area of the primary visual cortex, along with a rearrangement of neuronal connectivity, among other changes ^23^. We performed eyelid suture immediately after 14 days of weekly imaging under normal vision (3 sessions one week apart), followed by another two weeks of imaging under MD (imaging at 4d, 7d and 14d of MD), and studied the longitudinal distribution of myelin segments in single axons (Figure 3A). We imaged 39 mice for L2/3 CPN (Figure 3B), and 6 mice for L2/3 PV-INs (Figure 3F). For each animal, we examined the number of dynamic events and their length changes, for both pre-existing myelin sheaths and new myelin segments.

**Figure 3.**
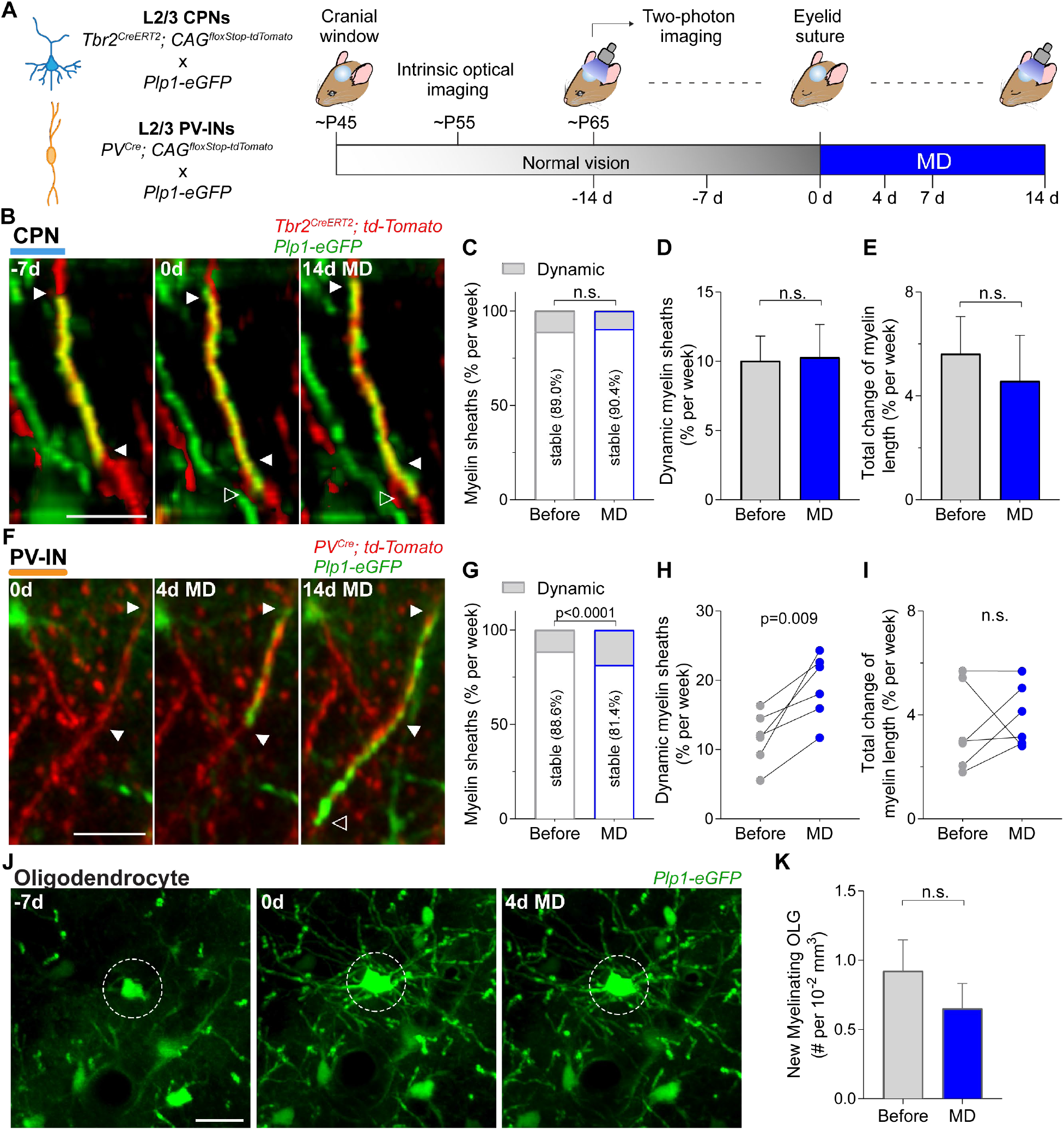
Experience-dependent remodeling of myelination profiles is neuron class-specific. (**A**) Schematic of cell-specific and myelin labeling strategy (left), and experimental time course (right). Cells in V1b were imaged at each indicated time point (bottom of the schematic). Monocular deprivation (MD) was induced by eyelid suture of the eye contralateral to the cranial window. (**B**) Representative three-dimensional images showing the elongation of a CPN myelin sheath (arrowheads) during normal visual experience. (**C**) Fraction of stable and dynamic myelin sheaths on CPNs (combined data from all mice). The time points were combined into two groups, before (−14d to −7d and −7d to 0d) and MD (0d to 7d and 7d to 14d). (**D**) Rate of myelin sheath dynamics for CPNs. Each mouse was analyzed as an independent statistical replicate (n = 39 mice). (**E**) Rate of change in internode length elicited by myelin remodeling. (**F**) MZP (5 frames) displaying the generation of a new myelin sheath on a PV-IN and its elongation during MD. (**G** to **I**) Analysis of PV-INs myelination (n = 6 mice), corresponding to (C) to (E). (**J**) MZP (22 frames) of a new myelinating oligodendrocyte (circle, 0 d). The circled cell at −7d is likely an oligodendrocyte progenitor cell. (**K**) Number of new myelinating oligodendrocytes generated before and during MD, normalized to the imaged volume (approximately 1.10^-2^ mm^3^ on average) (n = 19 mice). Scale bars: 10 μm, (B) and (F); 20 μm, (J). Data are mean ± s.e.m. For statistics, see table S1. n.s., not significant.

First, we compared the weekly rate of sheath dynamics on L2/3 CPNs during the two weeks of normal vision and the subsequent two weeks of MD (Figure 3C and 3D), and we found that the rate of remodeling remained unchanged (before, 10.0 ± 1.8 % per week vs. MD, 10.3 ± 2.4 % per week; paired t-test, p = 0.933; n = 39 mice, 146 sheaths). Similarly, MD did not affect the cumulative change in internode length caused by the population of dynamic sheaths (% of total length per week: before, 5.6 ± 1.4 % vs. MD, 4.6 ± 1.8 %; paired t-test, p = 0.647; Figure 3E). We measured the rate of dynamic sheaths and the normalized cumulative change in myelin length for each category of remodeling (elongation, contraction, and *de novo* generation), and determined that MD did not alter either of those parameters for any of the remodeling categories (Figure S4A-S4G). These results indicate that monocular deprivation does not affect the plasticity of myelination profiles in L2/3 CPNs.

Next, we asked whether PV-INs also localized to L2/3 had similar myelin dynamics in response to MD as their CPN neighbors. In contrast to CPNs, we found that PV-INs showed an increased rate of myelin dynamics upon MD (Figure 3G and 3H), indicating that experience drives adaptive modification of their longitudinal myelination profiles. We observed an increase in the rate of dynamic myelin sheaths (before, 11.6 ± 1.6 % per week vs MD, 19.1 ± 1.9 % per week; paired t-test, p = 0.009; n = 6 mice, 1630 sheaths; Figure 3H), and in 5 out of 6 mice there was also an increase in the cumulative change in their length (Figure 3I). By imaging the adjacent monocular visual cortex, we found that the increase in myelin sheath dynamics in PV-INs is exclusive to the binocular visual cortex and is absent in monocular visual cortex (Figure S5A-S5C), indicating that the changes are associated with the ocular dominance shift induced by MD (Figure S7). These findings demonstrate that sensory experience drives adaptive remodeling of myelination profiles specifically in PV-expressing interneurons, while CPN myelination shows only a continuation of homeostatic plasticity.

Finally, since new oligodendrocytes account for a substantial fraction of myelin remodeling under normal vision (Figure 2), we investigated the rate of generation of new myelinating oligodendrocytes during MD (Figure 3J). The initial density of mature oligodendrocytes was 47.1 ± 1.6 cells per 10^-2^ mm^3^ (P65, n = 19 mice) and, later, we found that the generation rate of new myelinating oligodendrocytes did not change upon MD (new myelinating oligodendrocytes per 10^-2^ mm^3^: before, 0.92 ± 0.22 vs MD, 0.65 ± 0.18; Wilcoxon matched-pairs signed rank test, p = 0.542, n = 19 animals; Figure 3K), in either L1 or L2/3 (Figure S6A-S6D). Thus, sensory experience drives cell type-specific changes in the neuronal myelination profile, but does not alter the rate of generation of new myelinating oligodendrocytes. Furthermore, we did found no changes in the proliferation (Figure S6F and S6G) and apoptosis (Figure S6H and S6I) of oligodendrocytes precursor cells upon MD, and neither any case of elimination of mature oligodendrocyte, either before or during MD (data not shown). These data reveal an unprecedented level of specificity for myelin plasticity, demonstrating that experience-dependent adaptive remodeling of myelination profiles is neuron class-specific and is sufficient to achieve circuit reconfiguration without the need to resort to *de novo* oligodendrogenesis.

### Temporal dynamics of adaptive remodeling of PV^+^ interneurons

GABAergic neurons in L2/3 of adult mouse V1b have been shown to respond to MD with a robust ocular dominance shift ^19^ accompanied by changes in their dendritic arbors that first undergo retraction and subsequently elongate ^21^. Therefore, we asked whether the experience-dependent increase in myelin dynamics in PV-INs also follows a defined spatiotemporal pattern, similar to functional neuronal responses and dendritic rearrangements, or if it is a sustained and uniform effect (Figure 4A). We analyzed myelin dynamics over the time course of MD-induced ocular dominance plasticity, and found that PV-INs presented a two-phase sequence of myelin dynamics. Throughout the first week of MD, there is an acute increase in myelination, by elongation of pre-existing sheaths and the generation of new myelin segments (Figure 4B, 4C and S7). In particular, the number of elongations and their change in length presented almost three-fold and two-fold increases, respectively, in response to experience. In the second week of MD (7-14 d MD), the elongation rate returned to baseline levels, accompanied by a two-fold and threefold increase in contractions and in the cumulative change in length, respectively (Figure 4B, 4C and S7). Surprisingly, we also observed full elimination of myelin segments upon MD in PV-INs (Figure 4D). This extreme form of contraction was not seen in PV-INs during normal visual experience or in CPNs under any condition. Notably, we found that MD also caused the recruitment of pre-existing oligodendrocytes to produce new myelin sheaths on PV-INs (Figure 4E and S5D). These results show that monocular deprivation triggers a series of events that progressively modify the myelination profile of PV-INs, including the generation of new sheaths by pre-existing oligodendrocytes.

**Figure 4.**
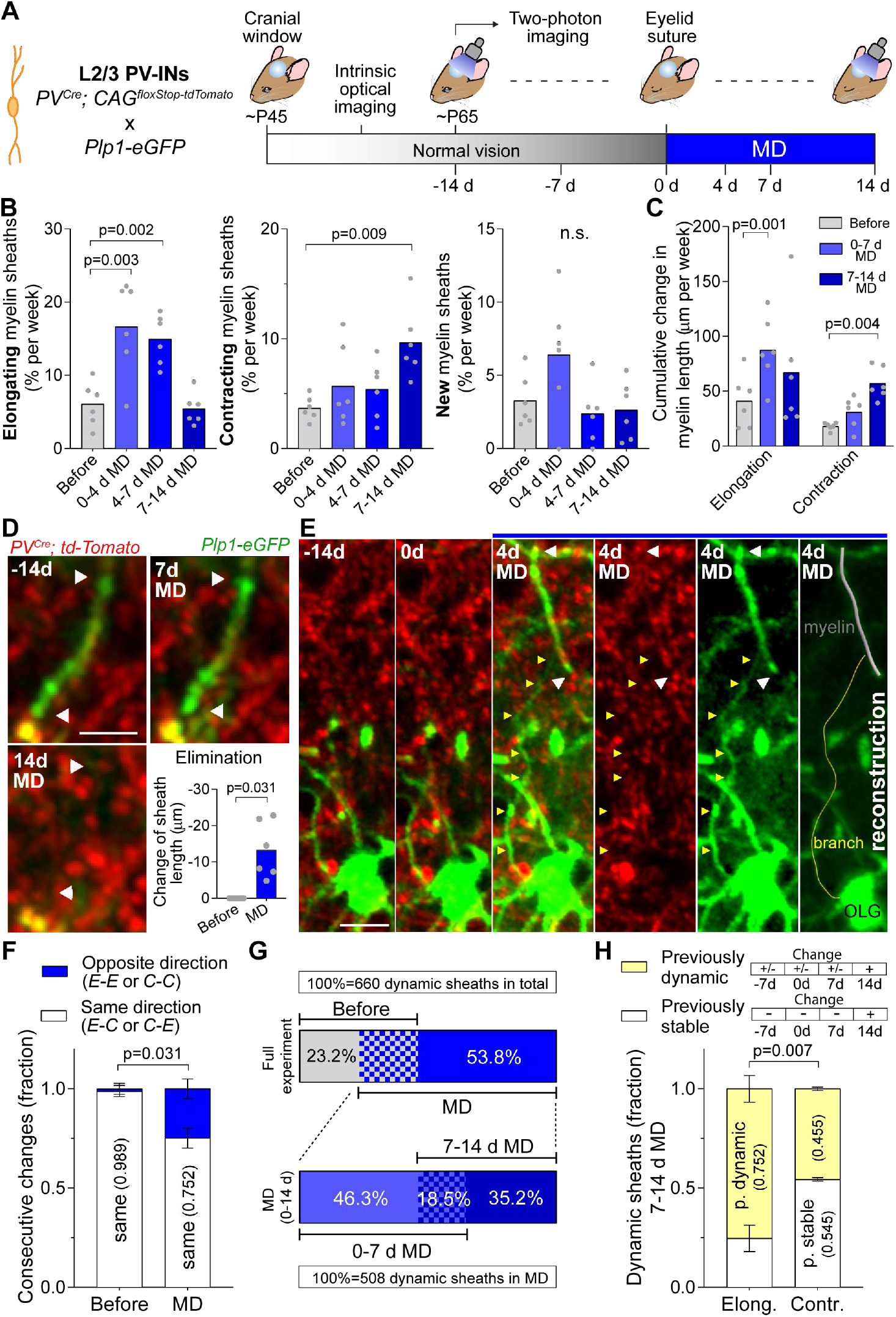
MD elicits temporally distinct phases of myelination dynamics in PV^+^ interneurons. (**A**) Schematic of PV-IN and myelin labeling strategy (left), and experimental time course (right). Cells in V1b were imaged at each indicated time point (bottom of the schematic). (**B**) Rate of myelin sheath elongation (left), contraction (center), and *de novo* generation (right) on PV-INs before and during MD. Each dot represents an animal. The data prior to MD (−14 d to 0 d) was combined into a single group (before). (**C**) Rate of absolute cumulative change in sheath length produced by contractions and elongations. (**D**) MZP (7 frames) displaying the elimination of a myelin sheath (arrowheads) upon MD, and total change in sheath length produced by full elimination of internodes (bottom right). (**E**) MZP (5 frames) showing a pre-existing oligodendrocyte (OLG, green) generating a new myelin sheath (white arrowheads) on a PV-IN (red) during MD. Yellow arrowheads indicate the branch that connects the myelin segment to the OLG cell body. A three-dimensional reconstruction is displayed at far right. (**F**) Fraction of consecutive changes in individual sheaths classified by the different combinations of direction changes: same (elongation/elongation or contraction/contraction) and opposite (elongation/contraction or contraction/elongation). (**G**) (Top) Percentage of myelin sheaths presenting changes only before MD (gray), only during MD (blue), and in both conditions (gridded, 23%). (Bottom) Percentage of myelin sheaths presenting changes only during the first week of MD (lighter blue), only the second week of MD (darker blue), and during both weeks of MD (gridded). (**H**) Fraction of myelin sheaths that experienced elongations or contractions during the second week of MD. For each type of remodeling, the myelin sheaths are classified by their previous plasticity: stable during the three weeks before (previously stable internode) or dynamic in any preceding session (previously dynamic internode). Scale bars: 5 μm, (D); 10 μm, (E). Data are mean ± s.e.m. For statistics, see table S1. n.s., not significant.

We next asked whether the myelin remodeling induced by MD represented an acceleration of the changes happening in basal conditions, or had different dynamics. Under normal vision, consecutive changes of a single myelin sheath length were always in the same direction; i.e., either successive elongations or successive contractions (Figure 4F). In contrast, during MD, 24.8 ± 2.0 % of successive changes on a single internode were in opposite directions (mostly elongation or *de novo* generation followed by contraction; Figure 4F), indicating that sensory experience induced an alteration in the pattern of myelin remodeling. We next asked whether the myelin sheaths that were remodeled during MD were the same as those that had been dynamic during normal vision. From the entire population of dynamic internodes identified during the full experiment, 53.8 % presented changes exclusively during MD while 23.2 % were dynamic solely under normal vision (Figure 4G), indicating that MD triggers the remodeling of a different set of sheaths rather than simply continuing changes in previously-dynamic sheaths. Moreover, a total of 81.5 % of sheaths dynamic during MD exhibited changes only during the elongation phase (0-7d MD, 46.3 %) or the retraction phase (7-14d MD, 35.2 %), but not both, indicating that the opposite phases affect two different subgroups of myelin segments (Figure 4G). In particular, 54.5 ± 0.8 % of contracting sheaths during the second week of MD had shown no change until then, while 75 ± 7 % of sheaths elongating during this period had shown previous changes (Figure 4H). These findings suggest that during the second phase of the myelin response to MD, a new subgroup of previously stable myelin segments are recruited for retraction.

Altogether, these results indicate that monocular deprivation drives the remodeling of myelin profiles in L2/3 PV-INs by triggering initial elongations in a set of myelin sheaths that had been until that point stable, followed by contraction in a different group of myelin sheaths. These data identify novel myelin dynamics triggered by monocular deprivation that distinguish the response to sensory perturbation from baseline homeostatic myelin plasticity.

### MD-induced remodeling of PV^+^ interneuron axons is independent of myelin plasticity

Since myelin placement is closely related to the structure and function of axons, we next interrogated the relationship between myelin plasticity and changes in axonal morphology and structure (Figure 5A and 5B). First, we investigated the impact of sensory experience on the nodes of Ranvier, short unmyelinated segments between myelin sheaths that are occupied by clusters of ion channels (Figure S2F). It has been shown that changes in myelin structure are accompanied by changes in the length of these nodes that modify the action potential propagation along the axon ^28^. We examined the dynamics of putative nodes of Ranvier in PV-INs (Figure 5A-5C), defined as short nodes (<5 μm) separating two neighboring myelin sheaths. Surprisingly, we found that putative nodes of Ranvier can be displaced along the axon during MD (10 out of 189 nodes in 6 mice; distance, 2.7 ± 0.3 μm), as a result of the contraction of one sheath and the elongation of another (Figure 5C and 5D). In contrast, node displacement was rarely seen under normal vision (2 out of 189 nodes). This result suggests that MD induces changes in the longitudinal pattern of PV-IN myelination and an associated increase in the rate of displacement of putative nodes of Ranvier.

**Figure 5.**
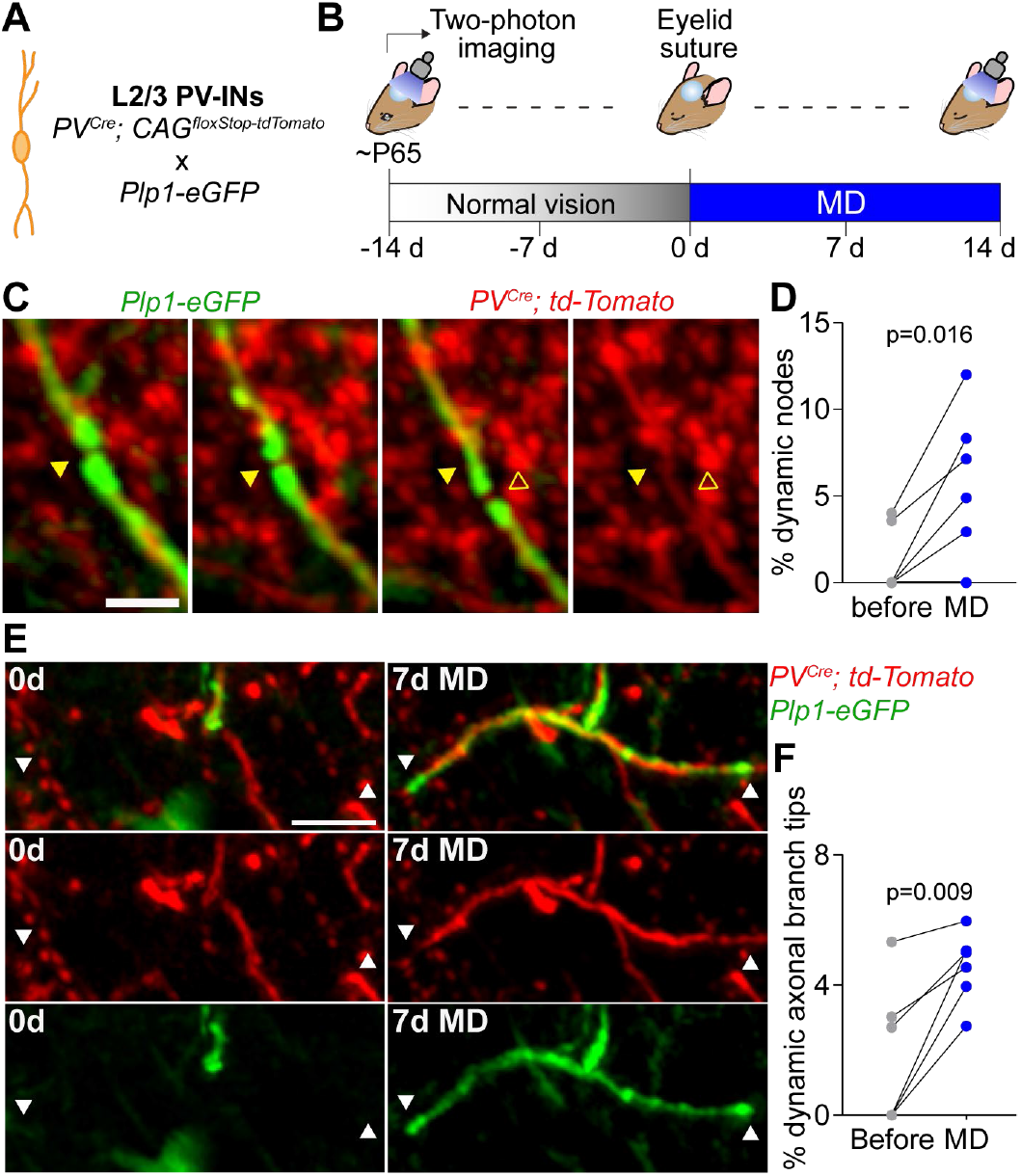
Sensory experience drives changes in the axonal arbor of PV^+^ interneurons. (**A**) Schematic of fluorescence labeling strategy. (**B**) Experimental time course. Cells in V1b were imaged at each indicated time point (bottom of the schematic). Monocular deprivation was induced by eyelid suture of the eye contralateral to the cranial window. (**C**) Single-frame images displaying a node in between two neighboring myelin sheaths, and a shift of its location during MD. Arrowheads indicate the putative node of Ranvier. (**D**) Number of relocated nodes of Ranvier before and during MD, normalized to the total number of nodes. (**E**) MZP (6 frames) showing a new myelin sheath (arrowheads) generated on a new axonal branch. (**F**) Number of dynamic axonal branch tips before and during MD, normalized to the total number of myelinated axons. Each dot represents an animal [(D) and (F)]. Scale bars: 5 μm, (C); 10 μm, (E). For statistics, see table S1.

Visual manipulations have been shown to induce reorganization of inhibitory neuron axons in adult visual cortex ^29^. We therefore examined axonal arbor remodeling in PV-INs, focusing on those branches that were myelinated at any point during the time course (Figure 5E). We found that myelin remodeling primarily occurred on axonal branches that did not exhibit changes in length during MD (95.6 ± 0.5 % of axons showing myelin remodeling, Figure 5F). This indicates that the experience-dependent remodeling of myelination profiles in PV-INs is not simply a downstream effect of axonal remodeling. In agreement with our findings that MD did not affect CPN myelin plasticity, we also did not observe axon remodeling in these neurons (data not shown). In the cases where axonal branch remodeling co-occurred with myelin plasticity in PV-INs, most of the combined axon and myelin remodeling events were observed when new sheaths were produced, although it remains possible that shorter contractions or elongations of myelin segments may have been excluded from the analysis due to higher uncertainty in their measurements (see Materials and Methods). Despite this technical limitation, we found that MD results in an increased number of dynamic axonal branch tips in PV-INs (Figure 5F), as well as an enlargement of axonal arbors in 5 out of 6 mice (change of axon length: normal vision, 19.3 ± 11.9 μm vs. MD, 47.89 ± 12.6 μm; paired t-test, p = 0.139, n = 6 mice). Our results indicate that PV-INs show independent increases in both axonal branch remodeling and myelination plasticity in response to monocular deprivation. Altogether, the data points at myelin and axon remodeling as pieces of a broader set of circuit adaptations orchestrated in response to perturbations in sensory experience.

## Discussion

The adult brain’s ability to change in response to experience arises from the coordinated modification of highly diverse neuronal and non-neuronal structures to ultimately modulate neuronal circuits. Even relatively small modifications in myelin sheath structure have the potential to substantially impact neural network function ^30^. Thus, understanding the dynamic mechanisms that govern myelin plasticity is a fundamental question in the neurosciences. Here we pioneer the investigation of cell type-specific experience-dependent myelin plasticity, and find that monocular deprivation drives an increase in myelin sheath remodeling in L2/3 PV^+^ interneurons, but does not alter the myelination dynamics of CPNs nor the generation of new myelinating oligodendrocytes. Prior reports have shown that neuronal activity and experience modulate myelination ^8,9,25,31,32^, and that active myelination by newly-formed oligodendrocytes is necessary for learning and memory ^12–14^ The results presented here suggests a novel framework for conceptualizing activity- and experience-dependent myelin plasticity, where cell type-specific adaptive remodeling of myelination allows different neurons to modify their individual functions and potentiates the tuning capacity of neuronal networks.

Our data shows that adaptive remodeling of axonal arbor and myelination profiles during MD is present in PV^+^ interneurons but not CPNs, and takes the form of an initial phase of myelin sheath elongations followed by a contraction phase. We hypothesize that a temporal series of sprouting and pruning might be a general strategy for myelin dynamics and circuit plasticity. Furthermore, our findings are in line with the more pronounced plasticity of L2/3 interneurons in response to MD, at both the physiological and the structural levels, compared to pyramidal cells ^19–22^, suggesting that sensory experience drives orchestrated sets of circuit adaptations specific to individual neuronal subtypes.

Prior studies have shown that oligodendrogenesis and the subsequent myelination is required for memory and learning ^12–14^. These studies, and work presented here show that new oligodendrocytes produce a large number of new myelin sheaths ^7,25^. However, our results demonstrate that MD does not stimulate the formation of new myelinating oligodendrocytes. Rather, it induces remodeling of pre-existing myelin sheaths on PV^+^ interneurons. We speculate that the reconfiguration of network connectivity that underlies a functional ocular dominance shift likely requires a fine and precise tuning of individual myelination profiles instead of a broad addition of myelin. In line with this hypothesis, we observed recruitment of pre-existing oligodendrocytes to generate new myelin sheaths. Recently, it has been shown that surviving oligodendrocytes can establish new myelin segments in a cuprizone-induced model of demyelination ^25^, but our data now indicates that new sheaths in adult animals are not only produced by newly-formed but also by pre-existing oligodendrocytes in physiological conditions.

Restricting adaptive myelin modification to inhibitory PV^+^ interneurons vs. excitatory CPNs might be an adaptation to dynamically regulate network activity patterns, and, as a result, sensory information processing. Such control of network activity can be achieved by the tuning of inhibitory signal propagation as well as the synchronization of pyramidal cell ensembles and generation of network oscillations ^24,33^. We found that MD induces both myelin segment contractions as well as full elimination of myelin sheaths on PV^+^ interneurons. Our data therefore support the idea that functional optimization does not maintain maximally conductive axons; instead, precise and axon-specific regulation of myelination parameters is required for efficient circuit rearrangement ^30,34,35^.

Our data shows that CPN myelination remodeling appears to occur continuously in adult animals and is not affected by MD, suggesting that CPN myelination serves a role outside adaptations to sustained sensory changes. PV^+^ interneurons and CPNs have different temporal and spatial myelination profiles: in PV^+^ interneurons, the onset of and subsequent rapid increase in myelination takes place in juvenile animals ^17^, followed by a balanced remodeling of pre-existing sheaths in adulthood (Figure 1). In contrast, CPNs present a delayed onset as well as an extremely slow accumulation (Figure S3), supported by our finding that pre-existing myelin sheaths of young-adult mice display a clear bias for elongation (Figure 1). Accordingly, CPN neuronal adaptations in the short- and mid-term of the mouse lifecycle may not rely on CPN myelin modifications. Further experiments are therefore needed to elucidate the functional role of myelin in CPNs at different time scales.

Neuronal activity can regulate myelination ^8,9,31^, which in turn affects nervous system function. Given that myelin affects network activity through its close partner, the neuron, experience-dependent myelin plasticity and its bidirectional relationship with neuronal function must be studied in a cell type-specific context. Elucidating the molecular and cellular mechanisms underlying cell type-specific myelination is a key step for glia research. Our data demonstrate cell type-specific dynamics of myelination plasticity, even when the distinct neuronal subpopulations are interconnected within the same circuit, surrounded by a shared environment, and myelinated by a common set of oligodendrocytes. The cell type-specificity of myelin plasticity observed here could potentially be related to the bias for inhibitory axons displayed by a subset of the oligodendrocytes in V1 ^17^, or to a more general heterogeneity in the glia population ^36^. An alternative, but not mutually exclusive, explanation might be that transcriptionally-distinct classes of neurons display different molecular signatures, such as cell surface markers, to individualize interactions with oligodendrocytes. The immense diversity of cell types in the mammalian neocortex is a key element for its higher-order functions, and it is likely that differential myelin plasticity plays a major role in the functional implementation of neuronal diversity.

## Materials and Methods

### Mice

We used the following genetically modified mouse lines: Plp1-eGFP (*37*), Pvalb^tm1(cre)Arbr^ (PV^Cre^) (*38*), Tbr2^CreERT2^ (unpublished, kindly provided by Josh Huang) and CAG^floxStop-tdTomato^ (Ai14) (B6;129S6-Gt(ROSA)26Sor^tm14(CAG-tdTomato)Hze^/J) conditional reporter line (*39*). Mice from these lines were crossed to generate *pV^Cre/+^; td-Tomato^+/-^; Plp1-eGFP^+/-^* mice and *Tbr2^CreERT2/+^; td-Tomato^+/-^; Plp1-eGFP^+/-^* mice to label oligodendrocytes along with parvalbumin-expressing interneurons (PV-IN) and callosal projection neurons (CPN), respectively. This fluorescence labeling strategy allowed us to visualize myelinating oligodendrocytes and myelin sheaths by eGFP, simultaneously with neuronal cell bodies and axons by tdTomato. To induce tdTomato expression in the Tbr2^CreERT2^ line, pregnant mice were injected at E18.5 with 4-hydroxytamoxifen in corn oil (single dose of 1 mg tamoxifen per kg body weight); that induction protocol was used for all experiments except Figure S1D and S1E. Mice were maintained on a C57BL/6J background, and both male and female mice were used for experiments. All procedures were designed to minimize animal suffering and approved by the Harvard University Institutional Animal Care and Use Committee and the Massachusetts Institute of Technology Committee on Animal Care, and performed in accordance with institutional and federal guidelines.

### Surgical procedure

To allow long-term visualization of *in vivo* neuronal morphology and myelination profiles, a 5 mm glass coverslip replacing a skull area (cranial window) was implanted over the right visual cortex of young adult mice (P42–P49) as previously described (*40*). Two to three weeks later, optically clear windows were selected for *in vivo* two-photon imaging. Animals were housed in groups of 2-4 mice per cage from weaning until the day before the first session of imaging (P60-P71); then, mice were singly housed for the remainder of the experiment. Sulfamethoxazole (1 mg ml^-1^) and trimethoprim (0.2 mg ml^-1^) were chronically administered in the drinking water through the final imaging session to maintain optical clarity of implanted windows.

### Optical intrinsic signal imaging

For functional identification of monocular and binocular visual cortex (V1b), optical imaging of intrinsic signal and data analysis were performed as described previously (*41*). Briefly, animals were mildly anesthetized with 0.75 % – 1 % isoflurane, restrained using a head mount, and placed facing a monitor. For visual stimuli, a horizontal bar (5° in height and 73° in width) drifting up with a period of 12 s was presented for 60 cycles on a high-refresh-rate screen positioned 25 cm in front of the animal. Images were acquired continuously under 610-nm illumination with an intrinsic imaging system (LongDaq Imager, Optical Imaging Inc.) through a 2.5X/0.075 NA objective (Zeiss). Cortical intrinsic signal was computed by extracting the Fourier component of light reflectance changes matched to stimulus frequency from 4×4 spatially binned images. The fractional change in reflectance represents response magnitude, and the magnitude maps were thresholded at 30 % of the peak-response amplitude to define a response region. Primary visual cortex was determined by stimulation of both eyes, while V1b was determined by stimulation of the ipsilateral eye. Monocular visual cortex was determined by subtracting the V1b map from the map of primary visual cortex.

### Two-photon imaging

After identifying the binocular visual cortex through optical intrinsic signal imaging and allowing sufficient time (about 3 weeks) for recovery from the cranial window surgery, adult mice were anesthetized (1 % - 1.25 % isoflurane) and head fixed in a stereotaxic frame for large-volume, high-resolution dual-color imaging using a custom-built two photon microscope. Since tdTomato and eGFP were used to label neurons and oligodendrocytes, respectively, the two fluorophores were simultaneously excited with a Mai Tai HP Ti:Sapphire laser (Spectra-Physics) set at 975 nm (CPN experiments) or 990 nm (PV-IN experiments) and pumped by a 14 W solid state laser delivering 100 fs pulses at a rate of 80 MHz. A 200×200×300 μm^3^ (CPN) or 200×200×200 μm^3^ (PV-IN) neuronal volume at 250 nm/pixel XY was acquired for each mouse by scanning the laser beams using galvanometric XY-scanning mirrors (6215H, Cambridge Technology). 0.8 μm/frame Z-resolution was achieved using a piezo actuator positioning system (Piezosystem, Jena). The output power from the 20X/1.0 NA water immersion objective (W Plan-Apochromat, Zeiss) was set to 50 mW. The emission signals were collected using the same objective, passed through an IR blocking filter (E700SP, Chroma Technology), and spectrally separated using a dichroic mirror at 560 nm. Emission signals were simultaneously collected with two independent photomultiplier tubes after passing through the appropriate bandpass filters (550/100 and 605/75). Then, two photon raw data were processed for spectral linear unmixing as described previously (*41*) and the images were converted to an RGB image z-stack using a home-built MATLAB script.

### Monocular deprivation

Monocular deprivation was performed by eyelid suture immediately after the third imaging session. Mice were anesthetized with 2 % isoflurane, lid margins were trimmed, and triple antibiotic ophthalmic ointment (Bausch & Lomb) was applied to the eye. Four to five individual stitches were placed using 6-0 vicryl along the extent of the trimmed lids. Suture integrity was inspected directly before each imaging session. Animals whose eyelids did not seal fully or had reopened were excluded from further experiments.

### Measurement of ocular dominance

Ocular dominance during normal condition and after monocular deprivation was determined from optical intrinsic signal images as previously described (*42*). The ocular dominance index (ODI) was calculated from the average of (C – I)/(C + I) for all pixels in the region identified as binocular visual cortex, where C and I represent the response magnitude of each pixel to the contralateral and ipsilateral eyes, respectively. The ODI ranges from +1 to −1, where a positive value indicates a contralateral bias and a negative value an ipsilateral bias.

### Analysis of fluorescence images

*In vivo* z-stacks collected from *PV^Cre/+^; td-Tomato^+/-^; Plp1-eGFP^+/-^* (to investigate myelination on PV-IN) and *Tbr2^CreERT2/+^; td-Tomato^+/-^; Plp1-eGFP^+/-^* (to study myelination on CPN) mice were acquired using two-photon microscopy. Data analysis was performed blind to the experimental conditions. Images were randomized (consecutive time points were paired for further comparisons) for analysis by blinded observers. To eliminate any hint about the experimental condition, the last imaging session was paired with the first one; the results from the latter comparison was used to validate the intermediate changes between consecutive sessions.

#### Identifying myelin sheaths along single axons

Image z-stacks and time-series were traced and analyzed using Neurolucida (MicroBrightField, Inc.) and FIJI/ImageJ. All analysis was performed on unprocessed images, and for presentation in figures, image brightness and contrast levels were adjusted for clarity. For CPN experiments, cell bodies and their primary axons were segmented using the voxel scooping method for a semi-automated tracing in Neurolucida. Then, the eGFP^+^ processes colocalizing to tdTomato^+^ axonal processes were also segmented, in both CPN and PV-IN experiments. eGFP^+^ processes were scored as myelin sheaths if they presented colocalization of at least 80% extension to a tdTomato^+^ axon, the centerlines of the eGFP^+^ process and tdTomato^+^ axon underneath were separated no further away than 2 pixels (0.5 μm), and the structures have a minimal average signal intensity of at least four times above shot noise background levels. Myelin paranodes were identified by increased eGFP fluorescence intensity (Figure 1). A gap in the fluorescence signal greater than 1.6 μm was used to designate a break in the myelination and classified as a different myelin sheath. Myelin sheaths with terminals that could be confidently identified across all imaging sessions, not extending beyond the imaging volume or obscured by blood vessels, were monitored and included to further analysis.

The depth of each myelin sheath was determined by its midpoint along the axial position, relative to the pial surface, and verified by the L1-L2/3 border at approximately 90 μm below the pia. The interphase between L1 and L2/3 was identified in the z-stacks by the high density of cell bodies in L2/3 and also through post hoc DAPI staining. L2/3 CPNs have axonal projections to deeper layers of the cortex as well as long projections to the contralateral hemisphere. Since the myelin analysis was restricted to L1-L3 of the ipsilateral cortex, the proportion of myelinated CPNs we have reported is probably a lower limit.

#### Analyzing myelin sheath dynamics

Changes in myelin sheath length were independently validated by two investigators, using the intersection of cellular processes as stable landmarks across images to set reference lines (changes in xy plane) or identify equivalent frames (changes in z direction). Double confirmation by two investigators was required for a change in length to be included for further analysis. For changes in myelin length with a perpendicular orientation to images stacks, we set an inferior threshold of 1.6 μm, while for changes with a lateral orientation the threshold was set at 1.25 μm. In addition, we discarded the changes in length < 2.0 μm that were followed by partial or complete reversion in the change, assuming them to be false positive generated by the intrinsic variability across imaging sessions over a stable sheath.

For each mouse, the percentage of myelin sheaths elongating or retracting between two successive imaging sessions, relative to the total sheath number of the previous imaging session, were defined as the rates of myelin sheath elongations and contractions, respectively. Contractions included both retractions of existing sheaths as well as the elimination of entire internodes (we observed only 8 cases of full elimination in total), while elongations and generation of new myelin sheaths were analyzed separately. Rate of sheath dynamics was defined as the sum of the rates of sheath elongations, contractions and *de novo* generations. For all imaging intervals, rates of sheath remodeling were normalized to a ‘% per week’ unit by calculating the percent of remodeling sheaths, multiplied by 7, divided by the number of days between imaging sessions. The effect in length was analyzed analogously to the rate, by computing the total change in length generated by sheath remodeling, relative to the total length of all myelin sheaths. When plotting total change in length by dynamics sheaths (Figure 2I, 3E, 3I and S5C) or changes in length due to elongations and contractions in the same plot (Figure 1K and 4C), we reported absolute values for reduction in length.

Confocal images to interrogate CPN myelination over time (Figure S3) were analyzed as described before for *in vivo* z-stacks.

#### Analyzing oligodendrogenesis

When we analyzed the generation of new myelinating oligodendrocytes, cells were followed in three dimensions using custom FIJI scripts by defining eGFP^+^ cell bodies at each time point and recording xyz coordinates. New eGFP^+^ cells were identified as myelinating oligodendrocytes when at least 20 new myelin sheaths were observed in close proximity (less than 60 μm) to the new oligodendrocyte soma (*7*).

#### Analyzing axonal arbor and putative nodes of Ranvier

Axonal arbor remodeling was study on PV-INs. Because of the high density of PV-IN processes in L2/3, we only studied the segments of axon that were myelinated at some point during the experiments and were confidently identifiable along their full length.

Putative nodes of Ranvier were identified by measuring the fluorescence intensity across the putative node; if the average intensity between adjacent myelin sheaths decreased below two times the shot noise background levels, and the length of the gap between eGFP+ processes was < 5 μm, the structure was considered a node (*7*). Displacement of a node was defined as a shift > 1.5 μm of the node center to an area previously occupied by an eGFP^+^ process.

### Fluorescence immunohistochemistry

Immunohistochemistry was performed at P28 and onward time points in transgenic mice, as indicated in each figure. Animals were anesthetized with tribromoethanol (Avertin) and transcardially perfused with 0.1 M PBS (phosphate buffered saline, pH 7.4) followed by 4% paraformaldehyde, as described previously (*43*). Cortical tissue was then post-fixed overnight in 4% paraformaldehyde, followed by 3 x 10-minutes washes in 0.1 M PBS. Serial coronal sections (40 μm thick) were cut using a Leica microtome (VT1000 S), collected in PBS with 0.02 % sodium azide and stored at 4°C. Free-floating sections were blocked for 1 hour at room temperature in blocking buffer (PBS with 0.02% sodium azide, 0.3% BSA, 0.3% Triton X-100, and 8% serum of the species corresponding to the secondary antibody), and then incubated overnight at 4°C in blocking buffer with the following primary antibodies: rat anti-MBP (monoclonal, 1:100, Millipore), rabbit anti-Parvalbumin (1:500, Swant), mouse anti-Rorβ (monoclonal, 1:100, Cosmo Bio), rabbit anti-Cux1 (polyclonal, 1:100, Santa Cruz Biotechnology), mouse anti-Satb2 (monoclonal, 1:50, Abcam), rabbit anti-GABA (polyclonal, 1:1000, Sigma-Aldrich), rabbit anti-Olig2 (1:100, IBL America), rabbit anti-s100β (polyclonal, 1:2000, Abcam), mouse anti-Caspr (monoclonal, 1:100, UC Davis/NIH NeuroMab), mouse anti-Pan-Nav1 (monoclonal, 1:100, UC Davis/NIH NeuroMab), goat anti-PDGFRα (polyclonal, 1:200, Novus), rabbit anti-Ki67 (polyclonal, 1:200, Abcam) and rabbit anti-cleaved-caspase-3 (polyclonal, 1:300, Cell Signaling). Secondary antibody (all 1:750, TermoFisher) labeling was performed at room temperature for 2 hours as follows: goat anti-rat IgG (H+L) Alexa Fluor 647, donkey anti-mouse IgG (H+L) Alexa Fluor 647, goat anti-rabbit IgG (H+L) Alexa Fluor 488, goat anti-mouse IgG (H+L) Alexa Fluor 647, donkey anti-rabbit IgG (H+L) Alexa Fluor 546 and donkey anti-goat IgG (H+L) Alexa Fluor 488. Sections were mounted using ProLong Gold (Invitrogen P36930). Confocal imaging was performed on a Zeiss LSM 700 microscope using three different lasers: FITC (488 nm laser line excitation; 522/35 emission filter), Cy3 (555 nm excitation; 583 emission), and Cy5 (639 nm excitation; 680/32 emission). The primary visual cortex was first identified using a 10x objective, and imaged using a 20x and 40x objectives with a z-step size of 0.99 μm and 0.47 μm, respectively.

### Data presentation and statistical analysis

Table S1 contains the statistical tests used to measure significance, the corresponding significance level (P value) and sample size. Normality was assessed using Shapiro-Wilk’s test at a p value of 0.05. When a data set did not satisfy normality criteria, nonparametric statistics were applied. Two-tailed Mann-Whitney U test was used for single comparisons, and two-tailed Wilcoxon matched-pairs signed rank test was used for paired values. For normal distributions, homoscedasticity was assessed using Bartlett’s test and F-test, at a p value of 0.05. For homogeneous variances, two-tailed t-test was used for single comparisons, repeated-measures one-way ANOVA followed by post hoc Dunnett’s test was used for statistical analysis of time course data, and two-way ANOVA was used for two-factor data sets. Paired t-test was used to compare paired data. In the only case were variances were not homogeneous, a t test with Welch’s correction was used. Two-tailed Fisher’s exact test (small sample size) or Chi-square test were used in the analysis of contingency tables. No statistical methods were used to predetermine sample sizes, but our sample sizes are similar to those reported in previous publications (*6, 7, 25*) and consistent with those used in the field. Unless otherwise specified, data are presented as mean ± s.e.m. (text and figures); when the number of statistical independent replicates is greater than 15 per condition, individual values are not plotted for clarity (instead, we display a summary plot). Statistical tests were performed using GraphPad Prism version 8.4.1 (GraphPad Software, Inc) or MATLAB 2018a (The MathWorks Inc., Natick, MA), and p < 0.05 was considered statistically significant.

## Acknowledgments

We thank J.R. Brown, P. Oyler-Castrillo, former and present members of the Arlotta laboratory and Nedivi laboratory for insightful discussions and editing of the manuscript, J. Huang and W. Macklin for sharing the Tbr2^CreERT2^ and the Plp1-eGFP mouse lines, respectively. J. Boivin for helpful advice on *in vivo* 2P-imaging, and V. Vuong for outstanding technical support and for help with image analysis.

## Author contributions

P.A., E.N. and S.M.Y. conceived the experiments. S.M.Y. performed all the experiments, with help from K.M. S.M.Y. analyzed data, with help from V.J. P.A., E.N. and S.M.Y. wrote the manuscript with contributions from all authors. All authors read and approved the final manuscript.

