## Supplementary Materials for "Neuron-class specific responses govern adaptive remodeling of myelination in the neocortex"

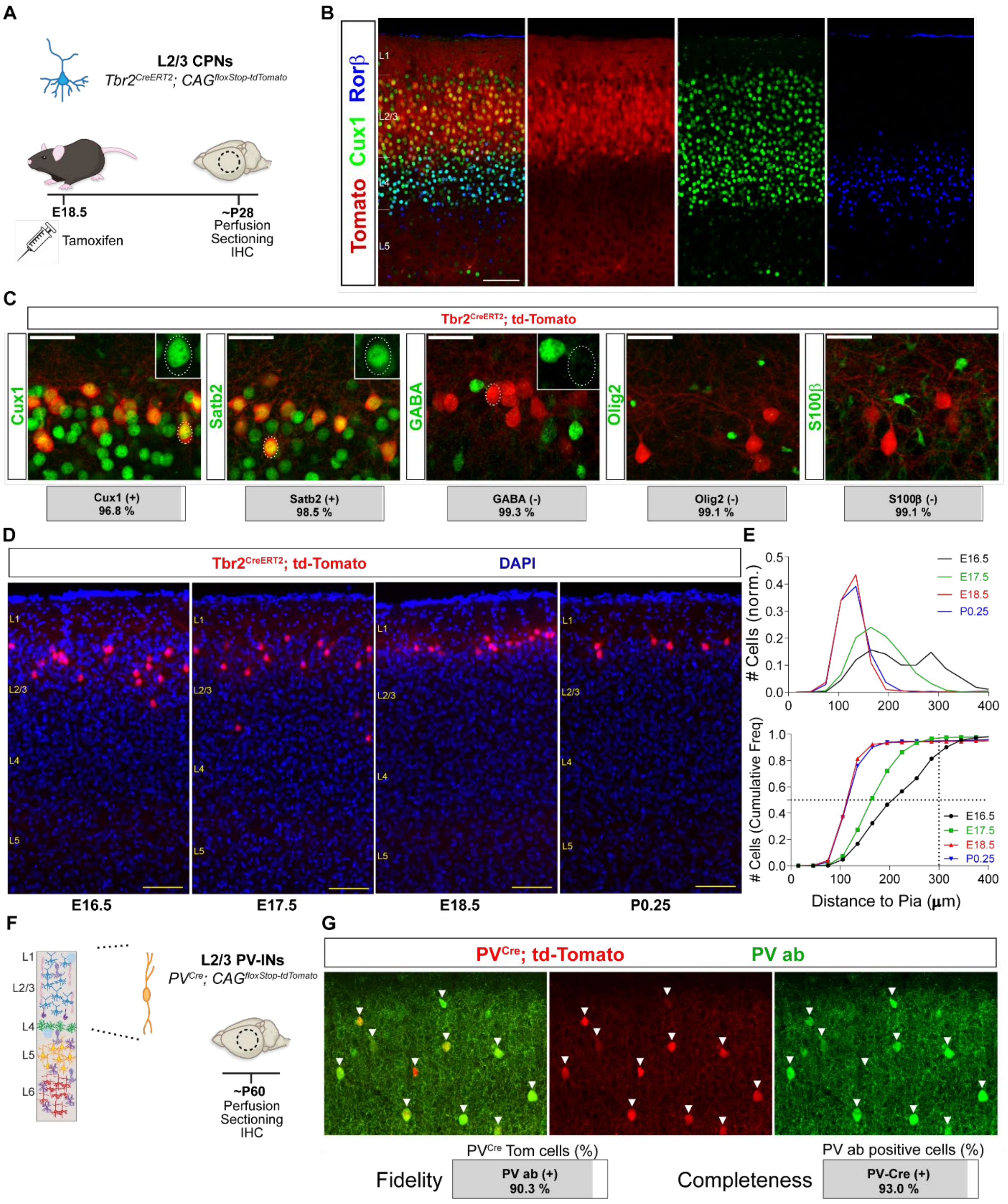


**Figure S1.**

**Characterization of Tbr2$^{Cre}$ and PV$^{Cre}$ transgenic lines.** (**A**) Schematic of experimental design for (B) to (E). (**B**) Immunohistochemistry for Cux1 and Rorβ in primary visual cortex of *Tbr2*$^{CreERT2/+}$*; td-Tomato*$^{+/-}$ mouse. (**C**) Immunohistochemistry for Cux1, Satb2, GABA, Olig2 and s100β in V1 L2/3 (n = 275-341 tdTomato$^{+}$ cells), with inset high magnification images of the indicated tdTomato$^{+}$ cells. Quantification is presented as percentage of tdTomato$^{+}$ cells (bottom). (**D**) Representative images showing the spatial distribution of labeled cells in V1 of *Tbr2*$^{CreERT2/+}$*; td-Tomato*$^{+/-}$ mice induced by tamoxifen administration at different ages (E16.5, E17.5, E18.5 and P0.25). (**E**) Distance from pial surface of tdTomato$^{+}$ cells in V1 of mice induced at different ages, expressed as fraction of total (top) and cumulative frequency distribution (bottom) (n = 6-8 mice and 660-1255 tdTomato$^{+}$ cells, per condition). (**F**) Schematic of experimental design for (G). (**G**) Immunohistochemistry for Parvalbumin (PV ab) in V1 L2/3 of *PV*$^{Cre/+}$*; td-Tomato*$^{+/-}$ mice. Arrows indicate tdTomato$^{+}$ cells. Quantification of the proportions of tdTomato$^{+}$ cells labelled by PV ab (left, fidelity) and PV-ab-labelled cells labelled with tdTomato (right, completeness) (n = 166 tdTomato$^{+}$ cells). Scale bars: 100 μm, (B); 20 μm, (C); 50 μm, (D) and (G).

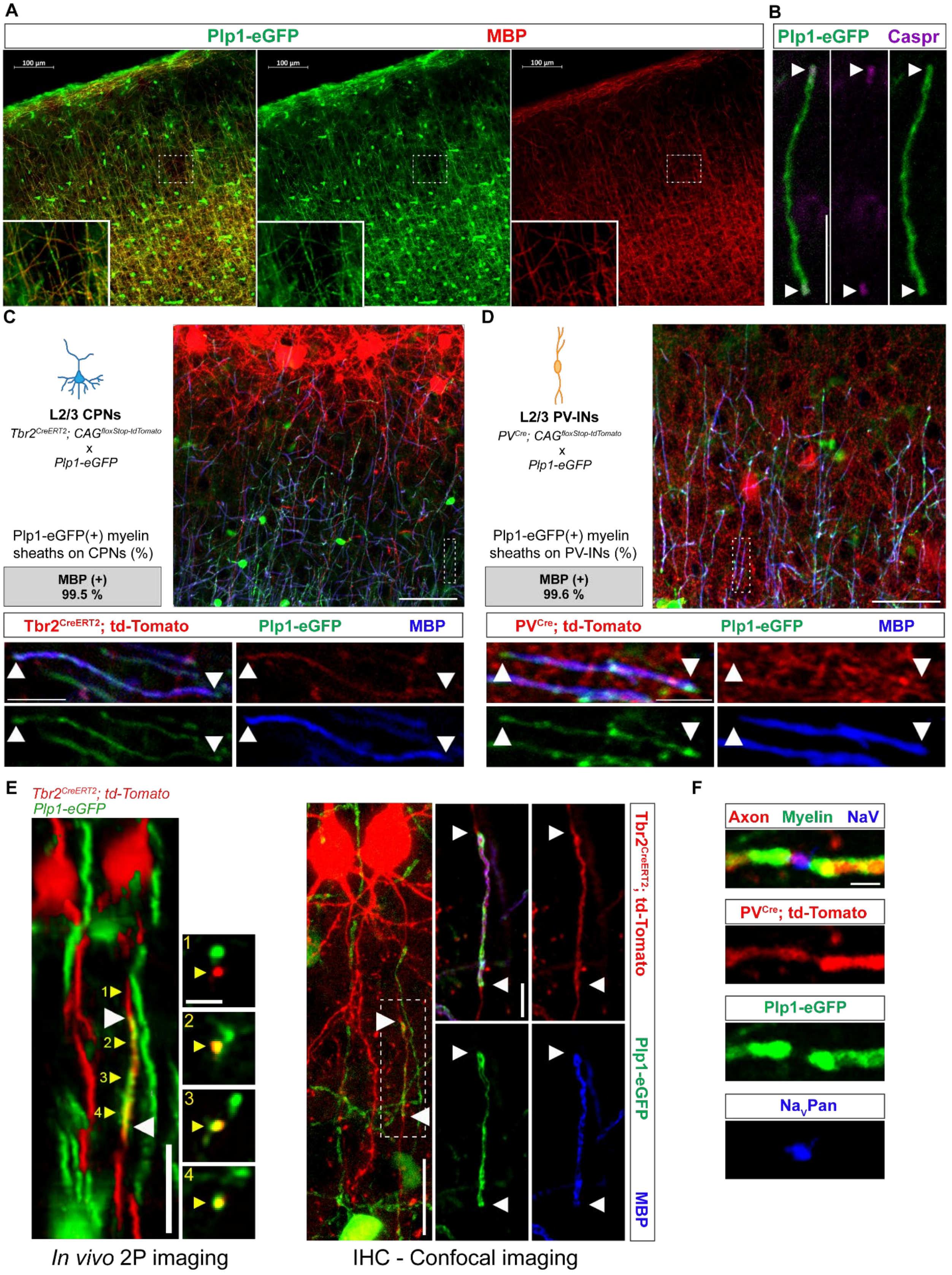

A
Plp1-eGFP MBP
100 µm
B
Plp1-eGFP Caspr
C
L2/3 CPNs
Tbr2CreERT2; CAGfloxStop-tdTomato
x
Plp1-eGFP
Plp1-eGFP(+) myelin sheaths on CPNs (%)
MBP (+)
99.5 %
Tbr2CreERT2; td-Tomato Plp1-eGFP MBP
D
L2/3 PV-INs
PVCre; CAGfloxStop-tdTomato
x
Plp1-eGFP
Plp1-eGFP(+) myelin sheaths on PV-INs (%)
MBP (+)
99.6 %
PVCre; td-Tomato Plp1-eGFP MBP
E
Tbr2CreERT2; td-Tomato
Plp1-eGFP
1
2
3
4
In vivo 2P imaging
Tbr2CreERT2; td-Tomato
Plp1-eGFP
MBP
IHC - Confocal imaging
F
Axon Myelin NaV
PVCre; td-Tomato
Plp1-eGFP
NaVPan

**Figure S2.**

**Characterization of Plp-eGFP transgenic line.** (**A**) Immunohistochemistry for MBP in P60 primary visual cortex of *Plp-eGFP*$^{+/-}$ mouse, with high-magnification insets of boxed areas. (**B**) Immunohistochemistry for Caspr in *Plp1-EGFP*$^{+/-}$ mouse. Arrows indicate the paranodes of an example myelin sheath. (**C** and **D**) Immunohistochemistry for MBP in V1 L2/3 of *Tbr2*$^{CreERT2/+}$*; td-Tomato*$^{+/-}$*; Plp-eGFP*$^{+/-}$ [(C), P210] and *PV*$^{Cre/+}$*; td-Tomato*$^{+/-}$*; Plp-eGFP*$^{+/-}$ [(D), P58] mice. For each panel: (Top left) Labeling strategy. (Top right) Example images of V1 L2/3. (Bottom) High magnification of boxed areas on top right, showing myelin sheaths (arrowheads) on single axons of a CPN (C) and a PV-IN (D); internodes express eGFP and MBP. (Center left) Percentage of eGFP$^{+}$ myelin sheaths on CPNs (C) and PV-INs (D) expressing MBP (CPN, n = 2 mice, 180 eGFP$^{+}$ sheaths; PV-IN, n = 2 mice, 233 eGFP$^{+}$ sheaths). (**E**) A single myelinated axon of a CPN was imaged by in vivo two-photon imaging (left; three-dimensional image) and after tissue fixation followed by immunohistochemistry for MBP (right). Cross-sections, at positions indicated by numbered yellow arrowheads, are displayed in the left panel. Single-frame images of the boxed area are presented in the right panel. (**F**) Immunohistochemistry for sodium channels (PanNa$_{V}$1) in *PV*$^{Cre/+}$*; td-Tomato*$^{+/-}$*; Plp-eGFP*$^{+/-}$ mouse. Scale bars: 100 μm, (A); 10 μm, (B), (C) inset and (D) inset; 50 μm, (C) and (D); 20 μm, (E); 5 μm, (E) inset; 2 μm, (F).

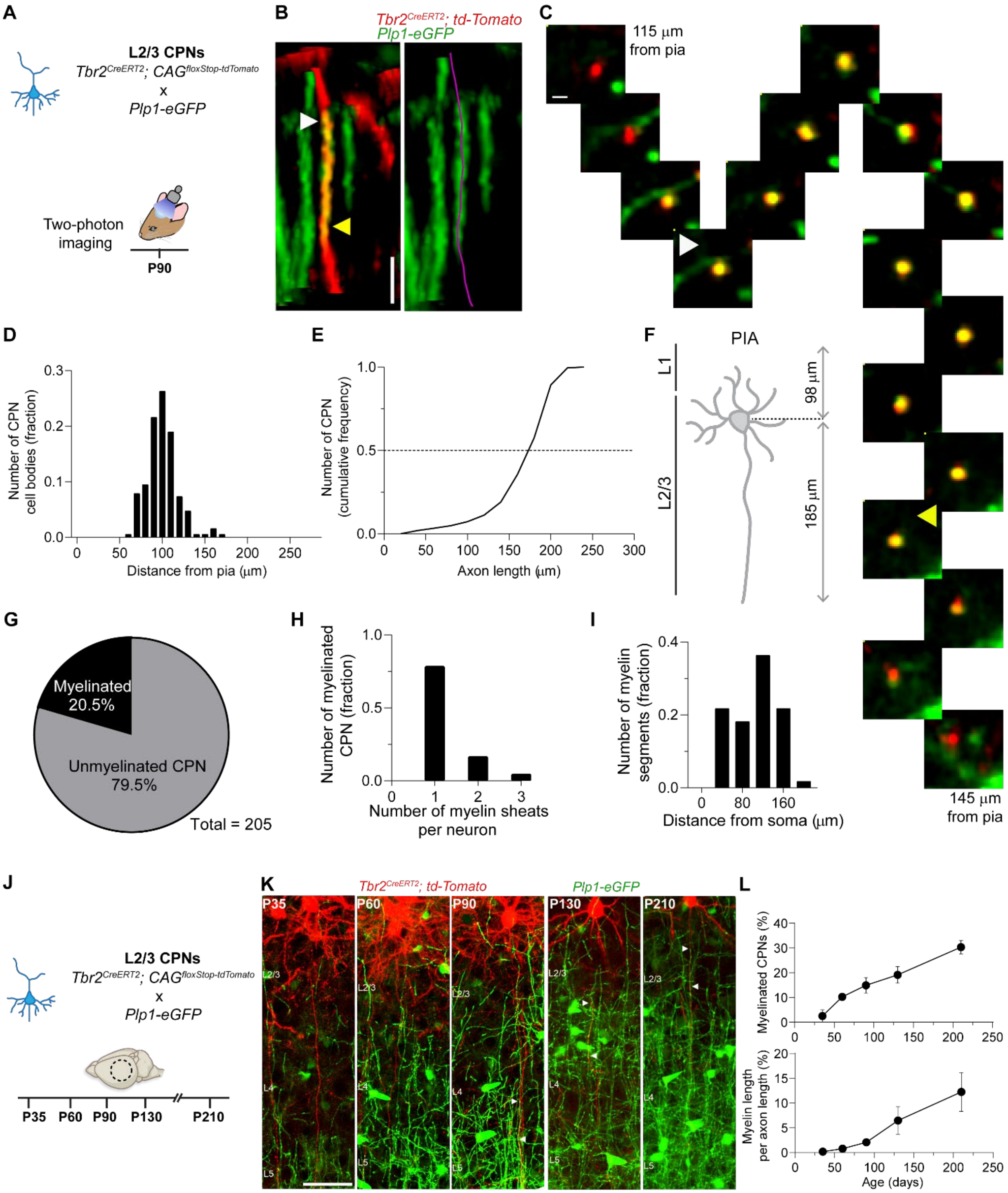


**Figure S3.**

**Myelination of L2/3 callosal projection neurons in adult mice.** (**A** to **I**) CPN myelination analyzed by *in vivo* two-photon imaging of P90 mice (related to Fig. 1 and 2). (**A**) Schematic of experimental design for (B) to (I). (**B**) Three-dimensional images showing a myelin sheath (arrowheads) on a CPN axon (reconstructed in purple, corresponding to Fig. 1C). (**C**) Sequence of singles frames displaying cross-sections of the myelinated axon presented in (B) (arrowheads correspond to the same positions). (**D** to **F**) Morphology of imaged CPN. Distance from pial surface of CPN somata (D), and primary axon length of CPNs (E), along with a schematic summary (F) (n = 19 mice, 205 cells). (**G**) Proportion of myelinated CPNs at P90. (**H**) Number of myelin sheaths per neuron (n = 56 sheaths). (**I**) Sheath position related to the soma. (**J** to **L**) CPN myelination in V1 over time. (**J**) Schematic of experimental design for (K) to (L). Brains were analyzed at each indicated time point. (**K**) Coronal sections showing increasing levels of myelination over time. Arrows indicate the tips of myelin sheaths on CPN. (**L**) Percentage of myelinated CPNs (top) and level of coverage (bottom) over time (n = 1-2 mice and 34-72 CPNs per time point). Scale bars: 10 μm, (B); 2 μm, (C); 50 μm, (K). Data are mean ± s.e.m.

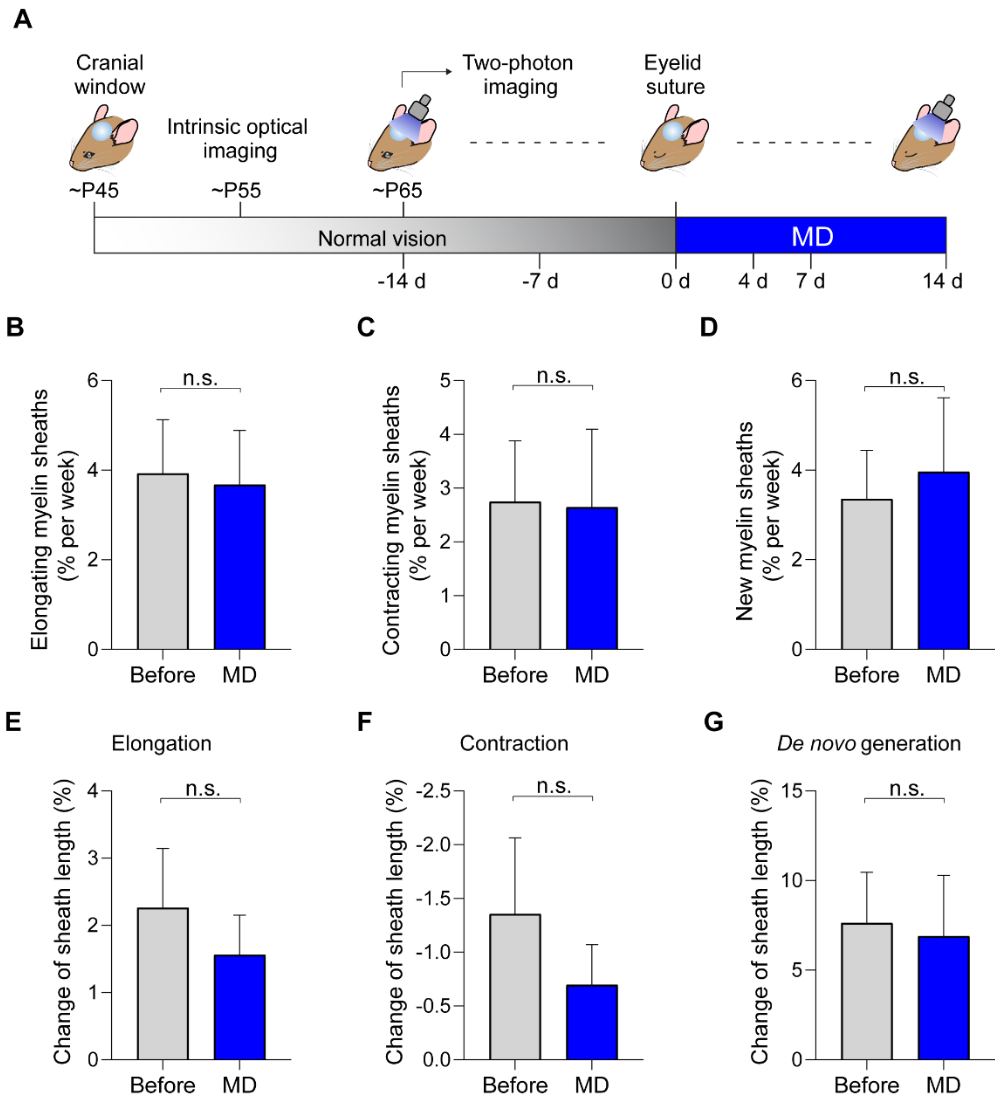


**Figure S4.**

**Monocular deprivation does not affect CPN myelin plasticity.** (**A**) Schematic of experimental design for (B) to (G). (**B** to **D**) Rate of myelin sheath remodeling before and during monocular deprivation (MD), analyzing elongation events (B), contraction events (C), and *de novo* generation (D) of myelin sheaths on L2/3 CPN. (**E** to **G**) Change in myelin segment length compared before and during MD, analyzing elongations (E), contractions (F), and *de novo* generation (G) of sheaths on CPN. Data are mean ± s.e.m. For statistics, see table S1. n.s., not significant.

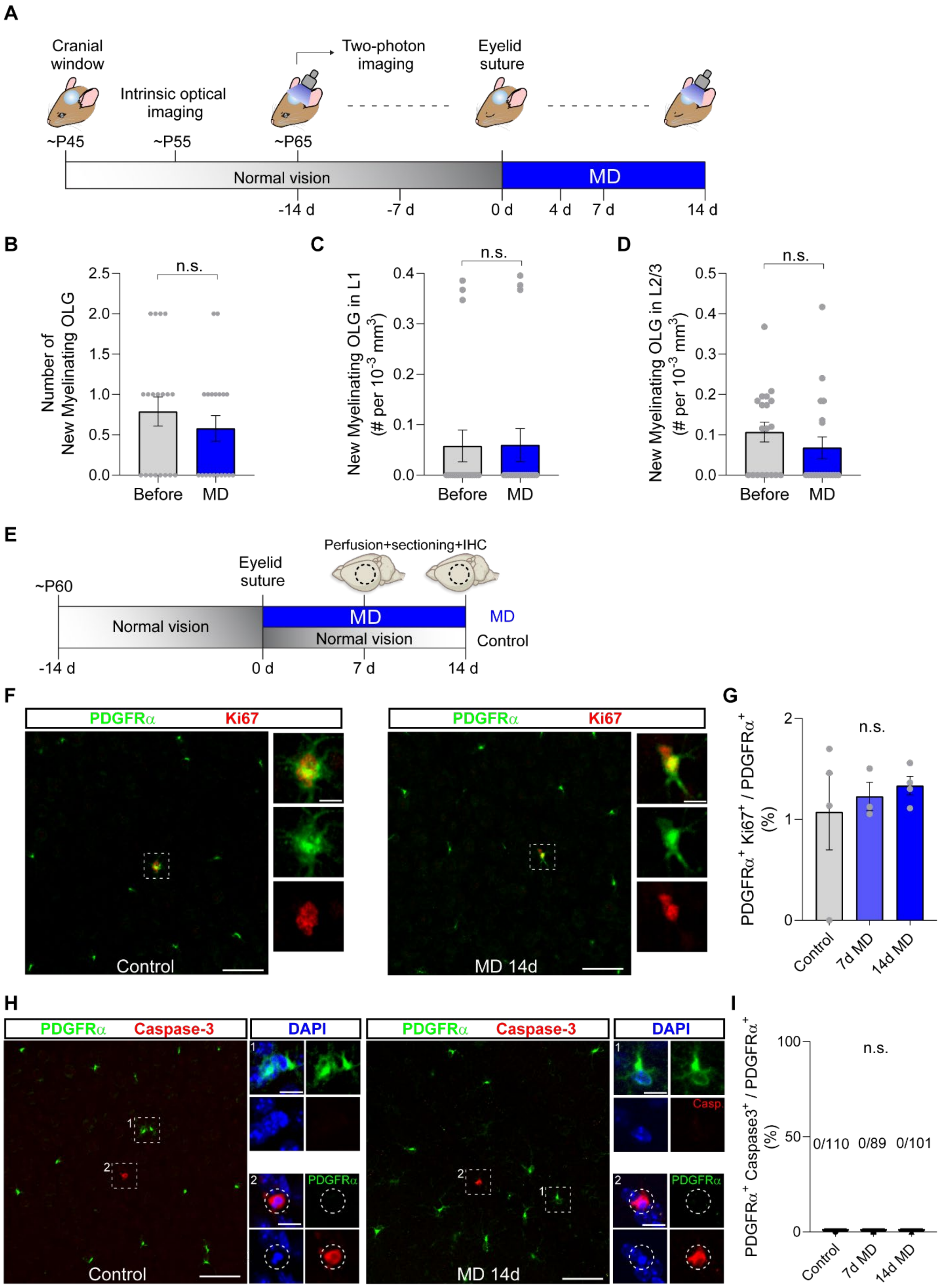

A
Cranial window
Intrinsic optical imaging
Two-photon imaging
Eyelid suture
~P45
~P55
~P65
Normal vision
MD
-14 d
-7 d
0 d
4 d
7 d
14 d
B
Number of New Myelinating OLG
n.s.
Before
MD
C
New Myelinating OLG in L1 (# per 10-3 mm3)
n.s.
Before
MD
D
New Myelinating OLG in L2/3 (# per 10-3 mm3)
n.s.
Before
MD
E
Perfusion+sectioning+IHC
Eyelid suture
~P60
Normal vision
MD
Normal vision
MD
Control
-14 d
0 d
7 d
14 d
F
PDGFRα Ki67
Control
PDGFRα Ki67
MD 14d
G
PDGFRα+ Ki67+ / PDGFRα+ (%)
n.s.
Control
7d MD
14d MD
H
PDGFRα Caspase-3
DAPI
PDGFRα
Control
PDGFRα Caspase-3
DAPI
Casp
PDGFRα
MD 14d
I
PDGFRα+ Caspase3+ / PDGFRα+ (%)
n.s.
0/110
0/89
0/101
Control
7d MD
14d MD

**Figure S5.**

**Monocular deprivation does not affect oligodendroglial lineage cells.** (**A** to **D**) Integration of new myelinating oligodendrocytes (OLG). (**A**) Schematic of experimental design for (B) to (D). (**B**) Number of new myelinating OLGs compared before and during MD. (**C** and **D**) Number of new myelinating OLGs generated before and during MD in L1 (C) and L2/3 (D), normalized to the imaged volume for each layer. (**E** to **I**) Dynamic of oligodendrocyte precursor cells (OPC). (**E**) Schematic of experimental design for (F) to (I). Tissue was collected 7 or 14 days after eyelid suture for the deprived group. (**F** to **I**) Proliferation (F and G) and apoptosis (H and I) of OPCs were studied by immunohistochemistry for Ki67 and cleaved caspase-3, respectively, while OPCs were stained by PDGFRα. Representative images are presented in (F) and (H), and percentage of PDGFRα-positive cells expressing each marker were plotted in (G) and (I). The number of double-positive cases are indicated in (I). Data are mean ± s.e.m. For statistics, see table S1. n.s., not significant.

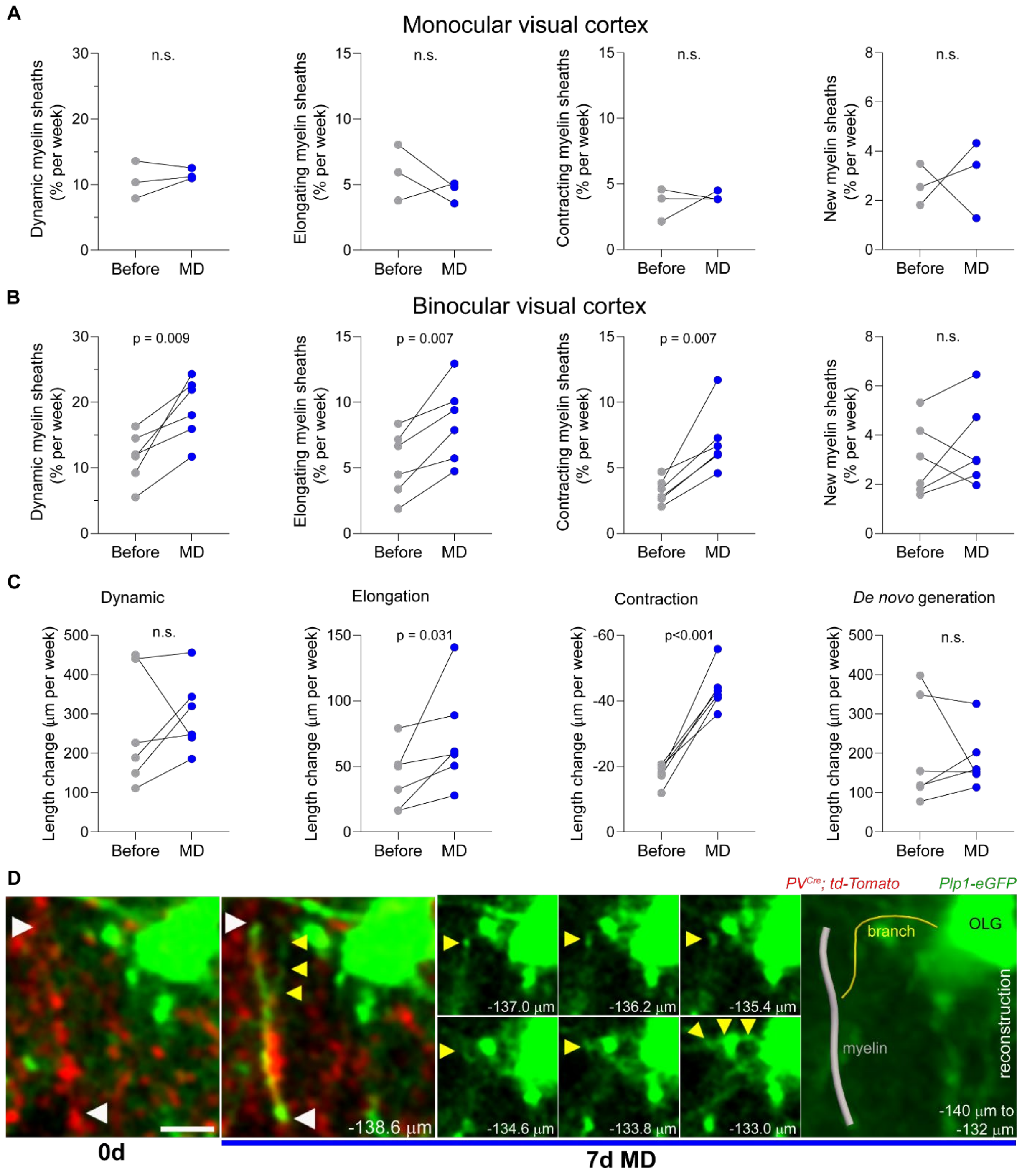


**Figure S6.**

**Monocular deprivation drives an increase of myelination dynamics in L2/3 PV$^{+}$ interneurons.** (**A** and **B**) Rate of myelin sheath remodeling before and during MD in monocular (A) and binocular (B) visual cortex. All dynamic sheaths on L2/3 PV-INs were analyzed both combined into a single group (left column), and as separated into 3 categories: elongation, contraction, and *de novo* generation. (**C**) Rate of change in myelin sheath length before and during MD in binocular visual cortex. (**D**) Single frame images showing a pre-existing oligodendrocyte generating a new myelin sheath on a PV-IN during MD. White and yellow triangles indicate the tips of the myelin sheath and the branch that connects it to the oligodendrocyte cell body (OLG), respectively. The distance from pia is indicated in each image. (Far right) Three-dimensional reconstruction displayed on a maximum z projection. Scale bar: 5 μm, (D). Each dot represents an animal. For statistics, see table S1. n.s., not significant.

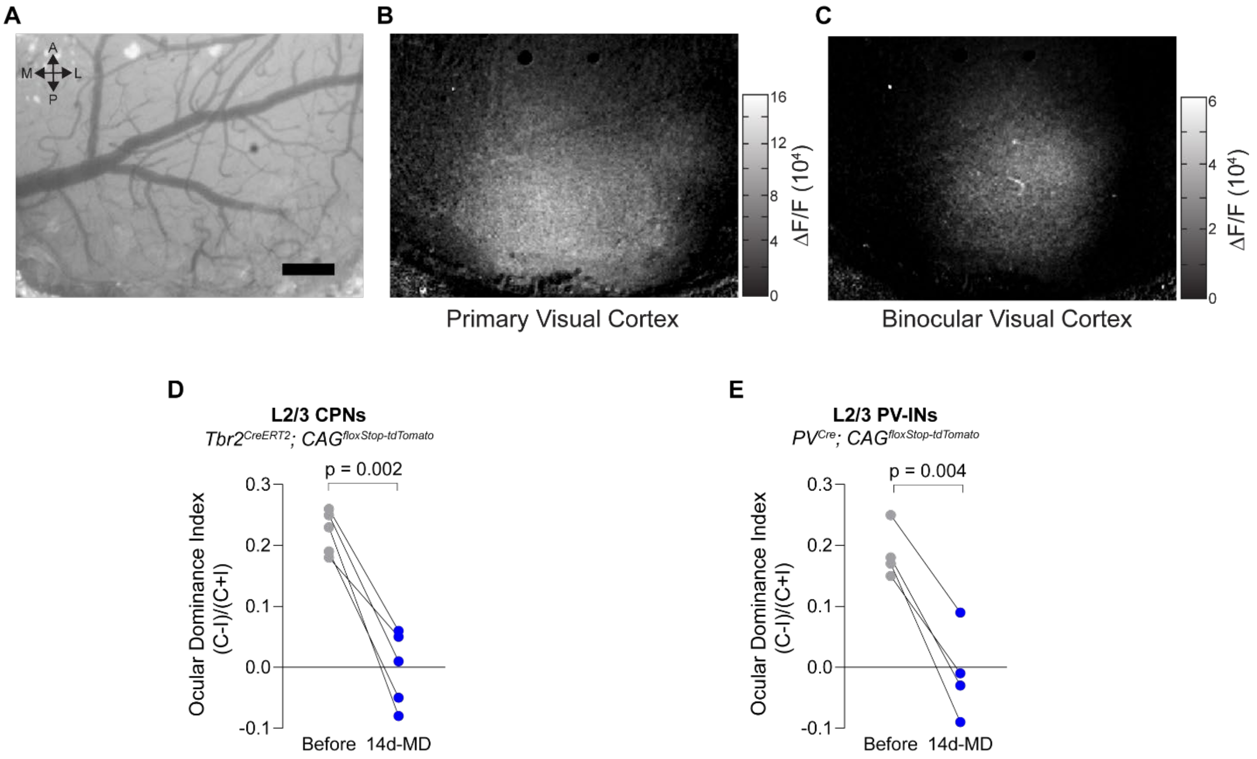


**Figure S7.**

**Ocular dominance plasticity through optical mapping of intrinsic signal.** (**A** to **C**) Imaging of an example animal. (**A**) Cortical blood vessel pattern of the region of interest. (**B** and **C**) Map of cortical responses measured as changes in reflectance upon visual stimulation to both eyes, identifying primary visual cortex (B), and the ipsilateral eye, identifying binocular visual cortex (C). (**D** and **E**) Ocular dominance index before and after 14-day monocular deprivation (14d-MD), for *Tbr2*$^{CreERT2}$*; td-Tomato* (D) and *PV*$^{Cre}$*; td-Tomato* mice (E). Each dot denotes an individual mouse. Scale bar: 200 µm, (A). For statistics, see table S1.

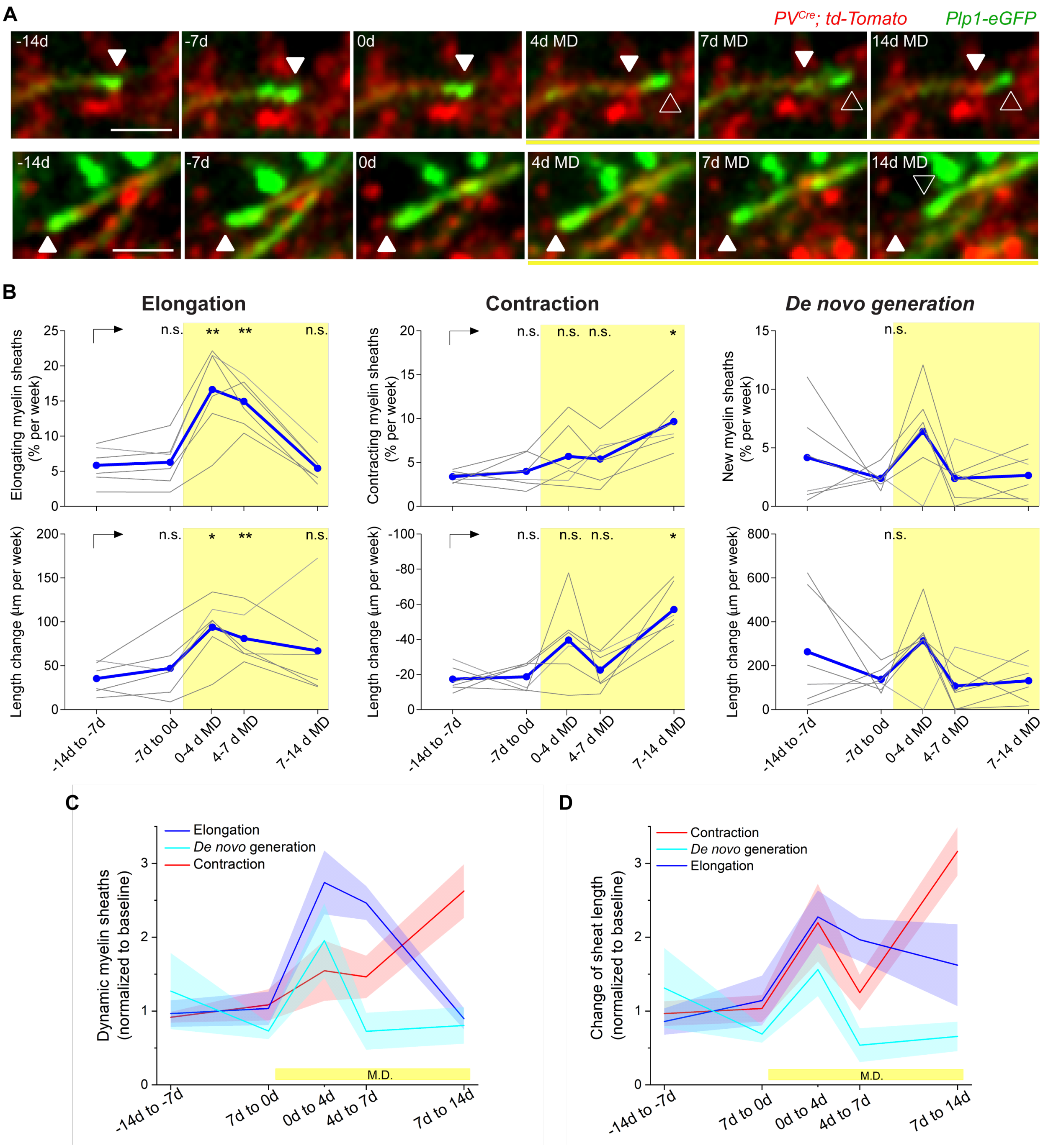


**Figure S8.**

**Monocular deprivation elicits a defined sequence of changes in PV-IN myelination.** (**A**) Single-frame images showing examples of early elongation (top) and late contraction (bottom) of myelin sheaths on PV-INs (arrowheads) during MD. (**B**) Rate of myelin sheaths dynamics (top row) and their length change (bottom row) in PV-INs imaged throughout a 14-d monocular deprivation time-course (yellow box). Remodeling events were categorized into elongation (left), contraction (center), and *de novo* generation (right). Individual animals are shown in gray and mean is represented in blue. (**C** and **D**) The data presented in (B) normalized to the baseline values (e.g., from -14 d to 0 d). Line and shaded area indicate mean and s.e.m., respectively. Scale bar: 5 μm, (A). For statistics, see table S1. n.s., not significant; * $p < 0.05$, ** $p < 0.01$.

**Table S1. Statistical analysis**

| Figure | Sample Size | Statistical Test | Values |
|---|---|---|---|
| **1G (bottom, length of myelin sheaths)** | CPN: 47 sheaths (from 20 mice)<br>PV-IN: 292 sheaths (from 3 mice) | Mann-Whitney U test (two-tailed) | $P < 0.0001$ |
| **1I** | CPN: 47 sheaths (from 20 mice)<br>PV-IN: 292 sheaths (from 3 mice) | Chi-square test | $P = 0.696$ |
| **1J** | CPN = 9 sheaths (from 20 mice)<br>PV-IN: 60 sheaths (from 3 mice) | Two-way ANOVA<br>factor one: neuron class<br>factor two: change type | Interaction: $p = 0.935$<br>$F(1,66) = 0.0068$<br>Neuron class: $p = 0.674$<br>$F(1,66) = 0.178$<br>Change type: $p = 0.208$<br>$F(1,66) = 1.616$ |
| **1K** | CPN: 20 mice<br>PV-IN: 3 mice | Wilcoxon matched-pairs signed rank test | $P = 0.039$ (CPN)<br>$P = 0.75$ (PV-IN) |
| **2C** | CPN: 56 sheaths (from 20 mice)<br>PV-IN: 380 sheaths (from 3 mice) | Fisher's exact test | $P = 0.302$ |
| **2D** | CPN: 56 sheaths (from 20 mice)<br>PV-IN: 380 sheaths (from 3 mice) | Mann-Whitney U test (two-tailed) | CPN: $p = 0.865$<br>PV-IN: $p = 0.051$ |
| **2F (top, # new myelin sheaths)** | 16 imaging sessions (from 4 mice) | Mann-Whitney U test (two-tailed) | $P = 0.002$ |
| **2F (bottom, total length of new myelin)** | 16 imaging sessions (from 4 mice) | Mann-Whitney U test (two-tailed) | $P = 0.004$ |
| **2H** | 551 pre-existing, 102 new +7d, and 38 new +14d sheaths (from 4 mice) | Fisher's exact test | $P = 0.001$ (pre-existing vs new +7d)<br>$P = 0.003$ (pre-existing vs new +14d) |

| | | | |
|---|---|---|---|
| **2I** | 44 pre-existing, 48 new +7d, and 9 new +14d, (from 4 mice) | One-way ANOVA followed by Dunnett's multiple comparison tests (against the pre-existing sheaths) | $P < 0.0001$<br>$F (2, 112) = 14.34$<br><br>$P < 0.0001$ (pre-existing vs new +7d)<br>P = 0.897 (pre-existing vs new +14d) |
| **3C** | 146 sheaths (in 451 neurons from 39 mice) | Fisher's exact test | P = 0.847 |
| **3D** | 39 mice | Paired t-test (two-tailed) | P = 0.933<br>($t_{38} = 0.0850$) |
| **3E** | 39 mice | Paired t-test (two-tailed) | P = 0.644<br>($t_{38} = 0.4654$) |
| **3G** | 1630 sheaths (from 6 mice) | Fisher's exact test | $P < 0.0001$ |
| **3H** | 6 mice | Paired t-test (two-tailed) | P = 0.009<br>($t_5 = 4.182$) |
| **3I** | 6 mice | Paired t-test (two-tailed) | P = 0.531<br>($t_5 = 0.6722$) |
| **3K** | 19 mice | Wilcoxon matched-pairs signed rank test (two-tailed) | P = 0.542 |
| **4B (left, elongating myelin sheaths)** | 6 mice | Repeated measures one-way ANOVA followed by Dunnett's multiple comparison tests (against before) | P = 0.0007<br>F (1.457, 7.283) = 26.78<br><br>P = 0.003 (before vs 0-4 d MD)<br>P = 0.002 (before vs 4-7 d MD)<br>P = 0.913 (before vs 7-14 d MD) |
| **4B (center, contracting myelin sheaths)** | 6 mice | Repeated measures one-way ANOVA followed by Dunnett's multiple comparison tests (against before) | P = 0.0025<br>F (2.593, 12.96) = 8.766<br><br>P = 0.415 (before vs 0-4 d MD)<br>P = 0.235 (before vs 4-7 d MD)<br>P = 0.009 (before vs 7-14 d MD) |
| **4B (right, new myelin sheaths)** | 6 mice | Repeated measures one-way ANOVA | P = 0.1018<br>F (1.309, 6.547) = 3.500 |

| | | | |
|---|---|---|---|
| **4C (elongation)** | 6 mice | Repeated measures one-way ANOVA followed by Dunnett's multiple comparison tests (against before) | P = 0.0398<br>F (1.102, 5.510) = 3.872<br><br>P = 0.001 (before vs 0-7 d MD)<br>P = 0.44 (before vs 7-14 d MD) |
| **4C (contraction)** | 6 mice | Repeated measures one-way ANOVA followed by Dunnett's multiple comparison tests (against before) | P = 0.0073<br>F (1.263, 6.316) = 13.82<br><br>P = 0.089 (before vs 0-7 d MD)<br>P = 0.004 (before vs 7-14 d MD) |
| **4D** | 6 mice (7 full eliminations) | Wilcoxon matched-pairs signed rank test (two-tailed) | P = 0.031 |
| **4F** | 6 mice (219 consecutive changes) | Wilcoxon matched-pairs signed rank test (two-tailed) | P = 0.031 |
| **4H** | 6 mice (232 dynamic sheaths) | Paired t-test (two-tailed) | P = 0.007<br>($t_5$ = 4.474) |
| **5D** | 6 mice (189 nodes) | Paired t-test (two-tailed) | P = 0.016<br>($t_5$=3.561) |
| **5F** | 6 mice (496 axonal branch tips) | Paired t-test (two-tailed) | P = 0.009<br>($t_5$ = 4.105) |
| **S1C** | 275 - 341 $tdTomato^+$ cells per condition | n/a | n/a |
| **S1E** | 660-1255 $tdTomato^+$ cells from 6-8 mice, per condition | n/a | n/a |
| **S1G** | 166 $tdTomato^+$ cells | n/a | n/a |
| **S2C** | 180 $eGFP^+$ sheaths from 2 mice | n/a | n/a |
| **S2D** | 233 $eGFP^+$ sheaths from 2 mice | n/a | n/a |
| **S3D** | 205 CPNs from 20 mice | n/a | n/a |
| **S3E** | 205 CPNs from 20 mice | n/a | n/a |

| | | | |
|---|---|---|---|
| **S3G** | 205 CPNs from 20 mice | n/a | n/a |
| **S3H** | 56 sheaths from 20 mice | n/a | n/a |
| **S3I** | 56 sheaths from 20 mice | n/a | n/a |
| **S3L** | 34-72 CPNs from 1-2 mice, per time point | n/a | n/a |
| **S4B** | 39 mice | Paired t-test (two-tailed) | P = 0.857 ($t_{38}$ = 0.1810) |
| **S4C** | 39 mice | Paired t-test (two-tailed) | P = 0.954 ($t_{38}$ = 0.0577) |
| **S4D** | 39 mice | Paired t-test (two-tailed) | P = 0.729 ($t_{38}$ = 0.3492) |
| **S4E** | 39 mice | Paired t-test (two-tailed) | P = 0.496 ($t_{38}$ = 0.6878) |
| **S4F** | 39 mice | Paired t-test (two-tailed) | P = 0.428 ($t_{38}$ = 0.8007) |
| **S4G** | 39 mice | Paired t-test (two-tailed) | P = 0.863 ($t_{38}$ = 0.1736) |
| **S5B** | 19 mice | Mann-Whitney U test (two-tailed) | P = 0.479 |
| **S5C** | 19 mice | Wilcoxon matched-pairs signed rank test (two-tailed) | P = 0.750 |
| **S5D** | 19 mice | Wilcoxon matched-pairs signed rank test (two-tailed) | P = 0.386 |
| **S5G** | 4 mice (control), 3 mice (7d MD), and 4 mice (14d MD) | Kruskal-Wallis test | P = 0.8497 |
| **S5I** | 4 mice (control), 3 mice (7d MD), and 4 mice (14d MD) | Kruskal-Wallis test | N/A |
| **S6A** | 3 mice (776 sheaths) | Wilcoxon matched-pairs signed rank test (two-tailed) | P = 0.750 (dynamic)<br>P = 0.500 (elongating)<br>P = 0.999 (contracting)<br>P = 0.750 (new) |

| | | | |
|---|---|---|---|
| **S6B** | 6 mice | Paired t-test (two-tailed) | P = 0.009 (dynamic)<br>($t_5$ = 4.182)<br>P = 0.007 (elongating)<br>($t_5$ = 4.480)<br>P = 0.007 (contracting)<br>($t_5$ = 4.430)<br>P = 0.403 (new)<br>($t_5$ = 0.9139) |
| **S6C** | 6 mice | Paired t-test (two-tailed) | P = 0.531 (dynamic)<br>($t_5$ = 0.6722)<br>P = 0.031 (elongating)<br>($t_5$ = 2.951)<br>P < 0.001 (contracting)<br>($t_5$ = 8.330)<br>P = 0.722 (new)<br>($t_5$ = 0.3764) |
| **S7D** | 5 mice | Paired t-test (two-tailed) | P = 0.002<br>($t_4$ = 7.612) |
| **S7E** | 4 mice | Paired t-test (two-tailed) | P = 0.004<br>($t_3$ = 8.251) |
| **S8B (top-left, % elongating sheaths)** | 6 mice | Repeated measures one-way ANOVA followed by Dunnett's multiple comparison tests (against the first time point) | P = 0.0003<br>F (1.683, 8.416) = 28.08<br><br>P = 0.818 (vs -7d to 0 d)<br>P = 0.004 (vs 0-4 d MD)<br>P = 0.001 (vs 4-7 d MD)<br>P = 0.977 (vs 7-14 d MD) |
| **S8B (top-center, % contracting sheaths)** | 6 mice | Repeated measures one-way ANOVA followed by Dunnett's multiple comparison tests (against the first time point) | P = 0.0023<br>F (2.771, 13.86) = 8.412<br><br>P = 0.838 (vs -7d to 0 d)<br>P = 0.489 (vs 0-4 d MD)<br>P = 0.358 (vs 4-7 d MD)<br>P = 0.019 (vs 7-14 d MD) |
| **S8B (top-right, % new sheaths)** | 6 mice | Repeated measures one-way ANOVA | P = 0.1456<br>F (1.928, 9.638) = 2.379 |

| | | | |
|---|---|---|---|
| **S8B (bottom-left, length change by elongation)** | 6 mice | Repeated measures one-way ANOVA followed by Dunnett's multiple comparison tests (against the first time point) | P = 0.0420<br>F (1.569, 7.845) = 5.200<br><br>P = 0.628 (vs -7d to 0 d)<br>P = 0.011 (vs 0-4 d MD)<br>P = 0.006 (vs 4-7 d MD)<br>P = 0.384 (vs 7-14 d MD) |
| **S8B (top-center, length change by contraction)** | 6 mice | Repeated measures one-way ANOVA followed by Dunnett's multiple comparison tests (against the first time point) | P = 0.0102<br>F (1.803, 9.017) = 8.272<br><br>P = 0.998 (vs -7d to 0 d)<br>P = 0.145 (vs 0-4 d MD)<br>P = 0.732 (vs 4-7 d MD)<br>P = 0.014 (vs 7-14 d MD) |
| **S8B (top-right, length change by *de novo* generation)** | 6 mice | Repeated measures one-way ANOVA | P = 0.1711<br>F (2.204, 11.02) = 2.065 |
| **S8C** | 6 mice | n/a | n/a |
| **S8D** | 6 mice | n/a | n/a |
